# Structures of complete extracellular receptor assemblies mediated by IL-12 and IL-23

**DOI:** 10.1101/2023.03.13.532366

**Authors:** Yehudi Bloch, Jan Felix, Romain Merceron, Mathias Provost, Royan Alipour Symakani, Robin De Backer, Elisabeth Lambert, Savvas N. Savvides

## Abstract

Cell-surface receptor complexes mediated by pro-inflammatory Interleukin-12 and -23, both validated therapeutic targets, are incompletely understood due to the lack of structural insights into their complete extracellular assemblies. Furthermore, there is a paucity of structural details describing the IL-12:receptor interaction interfaces, in contrast to IL23:receptor complexes. Here we report cryo-EM structures of fully assembled IL-12/IL-23:receptor complexes comprising the complete extracellular segments of the cognate receptors. The structures reveal important commonalities but also surprisingly diverse features. Whereas IL-12 and IL-23 both utilize a conspicuously presented aromatic residue on their α-subunit as a hotspot to interact with the N-terminal Ig-domain of their high affinity receptors, only IL-12 juxtaposes receptor domains proximal to the cell-membrane. Collectively, our findings will enable a cytokine-specific interrogation of IL-12 and IL-23 signaling in physiology and disease.

## Main

The interleukin 12 (IL-12) family of cytokines and cognate receptors are critically important in innate and adaptive immunity, mainly by regulating T-cell populations^1^. Among all cytokines, the IL-12 cytokine family is unique because it is defined by heterodimeric cytokines comprising permutable α- and β-subunits that activate combinations of specific and shared receptors to elicit signaling that can span the entire spectrum of pro-inflammatory and immunosuppressive outputs^2–4^. The two founding members of the family, IL-12 and IL-23, serve as pro-inflammatory and pro-stimulatory cytokines in the development of T_h_1 and T_h_17 subsets of helper T-cells, respectively^1,5^. Heterodimeric IL-12 and IL-23 share IL-12B (also, termed p40) as their β-subunit, which interacts with IL-12 receptor β1 (IL-12Rβ1)^6^, and are distinguished by their usage of their α-subunits IL-12A (p35) and IL-23A (p19)^7–9^, which interact with IL-12 receptor β2 (IL-12Rβ2) and IL23 receptor (IL-23R), respectively^6,10^ (Figure 1a). Although the field has been greatly enriched by diverse functional and structural studies^11–15^, essential insights into the architecture of the complete extracellular receptor complexes mediated by IL-12 and IL-23 have eluded the field. Here, we present structures of the extracellular receptor assemblies mediated by mouse IL-12 (mIL-12) and human IL-23 (hIL-23), as elucidated via cryo-electron microscopy (cryo-EM) to 3.6 Å and 3.5 Å resolution, respectively.

**Figure 1:**
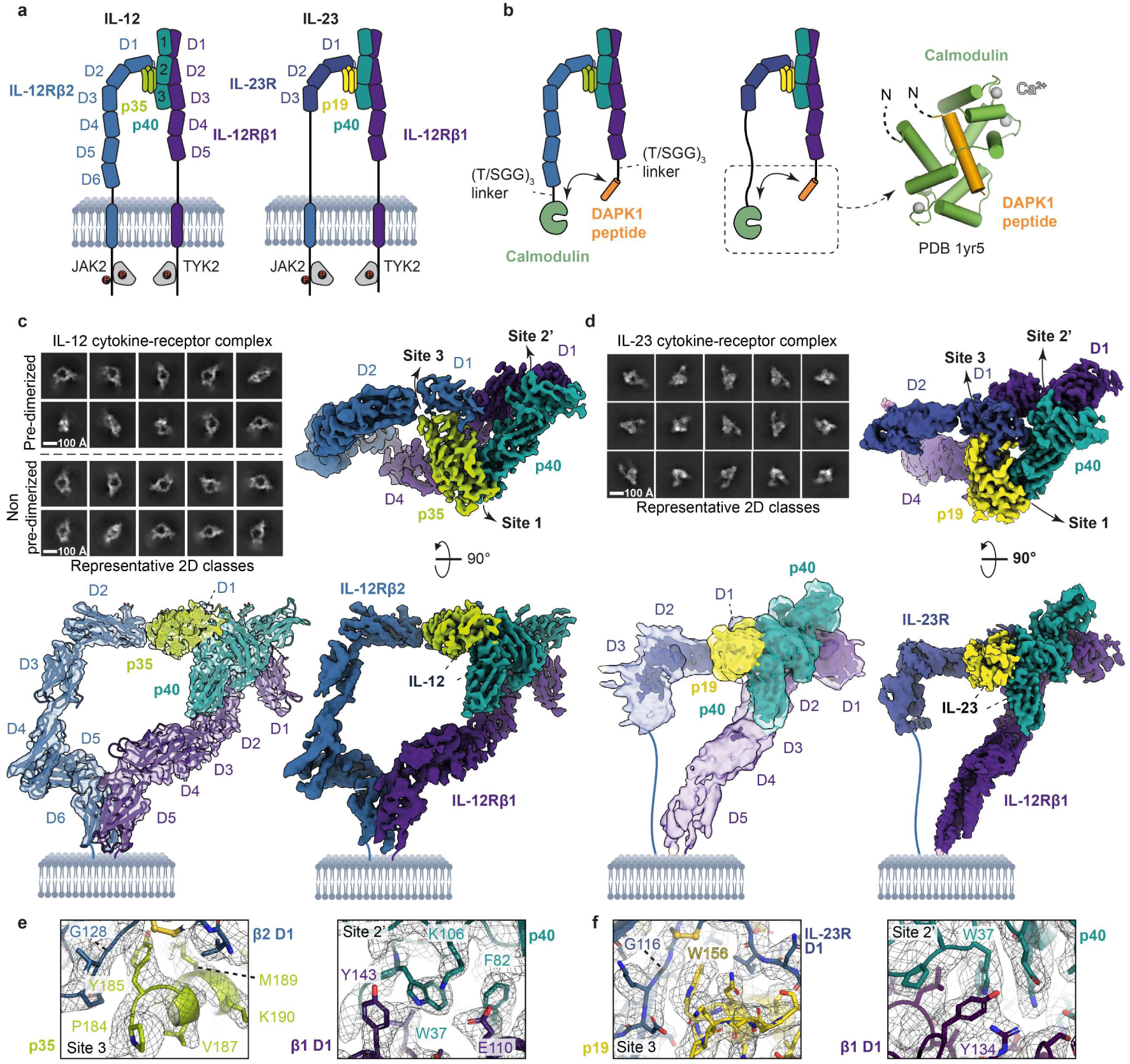
Structures of complete extracellular receptor assemblies mediated by IL-12 and IL-23. **a**, Schematic representation of IL-12 and IL-23 ligand-receptor assemblies on the cell surface (p35: IL-12A, p40: IL-12B, p19: IL-23A). **b**, Left: schematic representation of pre-dimerized extracellular IL-12 and IL-23 ligand-receptor assemblies. The calmodulin (CaM) and DAPK1 peptide (DAPK1_300-319_) complex structure (PDB code 1YR5) is shown on the right as a cartoon, with CaM in green, DAPK1_300-319_ in orange, and bound calcium (Ca^2+^) ions in grey **c**, Representative 2D class averages (top left) of Pre-dimerized and Non pre-dimerized murine IL-12 (mIL-12) extracellular ligand-receptor complexes are shown alongside a top view (top right) and side views (bottom) of the highest resolution 3D class of the pre-dimerized mIL-12 extracellular ligand-receptor complex. A fit with the atomic model is shown as a cartoon in a transparent map (c, bottom left). mIL-12A (p35), mIL-12B (p40), mIL-12Rβ1_D1-D5_-DAPK1_302-330_ and mIL-12Rβ2_D1-D6_-CaM are shown in green, cyan, purple and blue respectively. DAPK1_300-319_ and CaM tags are invisible in the 3D map due to flexibility of the linkers. The displayed 3D map was sharpened using DeepEMhancer^27^. **d**, Representative 2D class averages (top left) of human IL-23 (hIL-23) extracellular ligand-receptor complexes are shown alongside a top view (top right) and side views (bottom) of the 3D map of the hIL-23 extracellular ligand-receptor complex. The 3D maps displayed on the bottom left correspond to a sharpened map displayed at different threshold levels to show the presence/absence of signal for the membrane proximal receptor domains. The 3D maps displayed on the top right and bottom right correspond to a DeepEMhancer^27^ sharpened map. hIL-23A (p19), hIL-12B (p40), hIL-12Rβ1_D1-D5_-DAPK1_302-330_ and hIL-23R-CaM are shown in yellow, cyan, purple and dark blue respectively. DAPK1_300-319_ and CaM tags are invisible in the 3D map due to flexibility of the linkers. **e**, Insets showing key interactions in the mIL-12A (p35) – mIL-12Rβ2 (left) and mIL-12B (p40) – mIL-12Rβ1 (right) interfaces. Atomic models are displayed as a cartoon fitted in the corresponding 3D map displayed as a blue mesh. mIL-12A (p40), mIL-12B (p40), mIL-12Rβ1-DAPK1_302-330_ and mIL-12Rβ2-CaM are shown in yellow, cyan, purple and blue respectively. **f**, Insets showing key interactions in the hIL-23A (p19) – hIL-23R (left) and hIL-12B (p40) – hIL-12Rβ1 (right) interfaces. Atomic models are displayed as a cartoon fitted in the corresponding 3D map displayed as a blue mesh. hIL-23A (p19), hIL-12B (p40), hIL-12Rβ1-DAPK1_302-330_ and hIL-23R-CaM are shown in yellow, cyan, purple and dark blue respectively.

To facilitate the assembly of extracellular IL-12/IL-23:receptor complexes that under physiological conditions benefit from the dimensionality restrictions of their membrane-bound context, we designed a heterodimerization tag consisting of residues 300-319 of human Death-associated protein kinase 1 (DAPK1) and residues 5-149 of human Calmodulin-1 (CaM)^16^, fused via a GTGGSGGSGG linker to the C-termini of IL-12Rβ1 and IL-12Rβ2/IL-23R, respectively (Figure 1b, Methods). In addition, as isolation of stable receptor complexes mediated by hIL-12 proved challenging due to the tendency of human IL-12Rβ1 to dissociate from the complex, we focused on complexes mediated by mIL-12. This approach led to much improved yields of stoichiometrically stable mouse IL-12:IL-12Rβ2:IL-12Rβ1 complex, which we could rationalize by the ~10-fold higher affinity of mIL-12 for mIL-12Rβ1 than the human counterparts (Extended Data Figure 1, Methods, Source Data). Interestingly, the affinity of mIL-12 for mIL-12Rβ1 is similar to that of affinity-matured hIL-12 for hIL-12Rβ1 ^15^, which could be traced to the fortuitous conservation of amino acid positions important for binding (S129 and Y151 in mIL-12 versus Y109S & Q132L in affinity-matured hIL-12) (Extended Data Figure 1c).

Processing of cryo-EM data obtained for purified mIL-12/hIL-23:receptor complexes (Extended Data Figure 2,3; Methods) resulted in two distinct 3D reconstructions for the mIL-12:IL-12Rβ1:IL-12Rβ2 complex at 3.6 Å and 4.6 Å resolution (Figure 2a,b; Extended Data Figure 4a, Table 1), and one 3D reconstruction for the hIL-23:IL-12Rβ1:IL-23R at 3.5 Å (Figure 1d, Extended Data Figure 4b, Table 1). The obtained maps for the receptor complexes mediated by mIL-12 and hIL-23 displayed a considerable spread in estimated local resolution (Extended Data Figure 4), with the interfaces between the cytokines and receptors being most ordered and the membrane-proximal receptor extremities the least. Model building in these maps was aided by available X-ray structures of hIL-23 in complex with truncated receptors^14,15^, AlphaFold^17^ models of individual receptor ectodomains, and X-ray structures of unbound mIL-12, of mIL-12B_C197S_ in complex with mIL-12Rβ1_D1-D2_, and a hIL-12Rβ1_D3-D5_:Fab complex all determined herein at 2.9 Å, 2.2 Å and 2.6 Å resolution, respectively (Supplementary Table 1, Methods).

**Figure 2:**
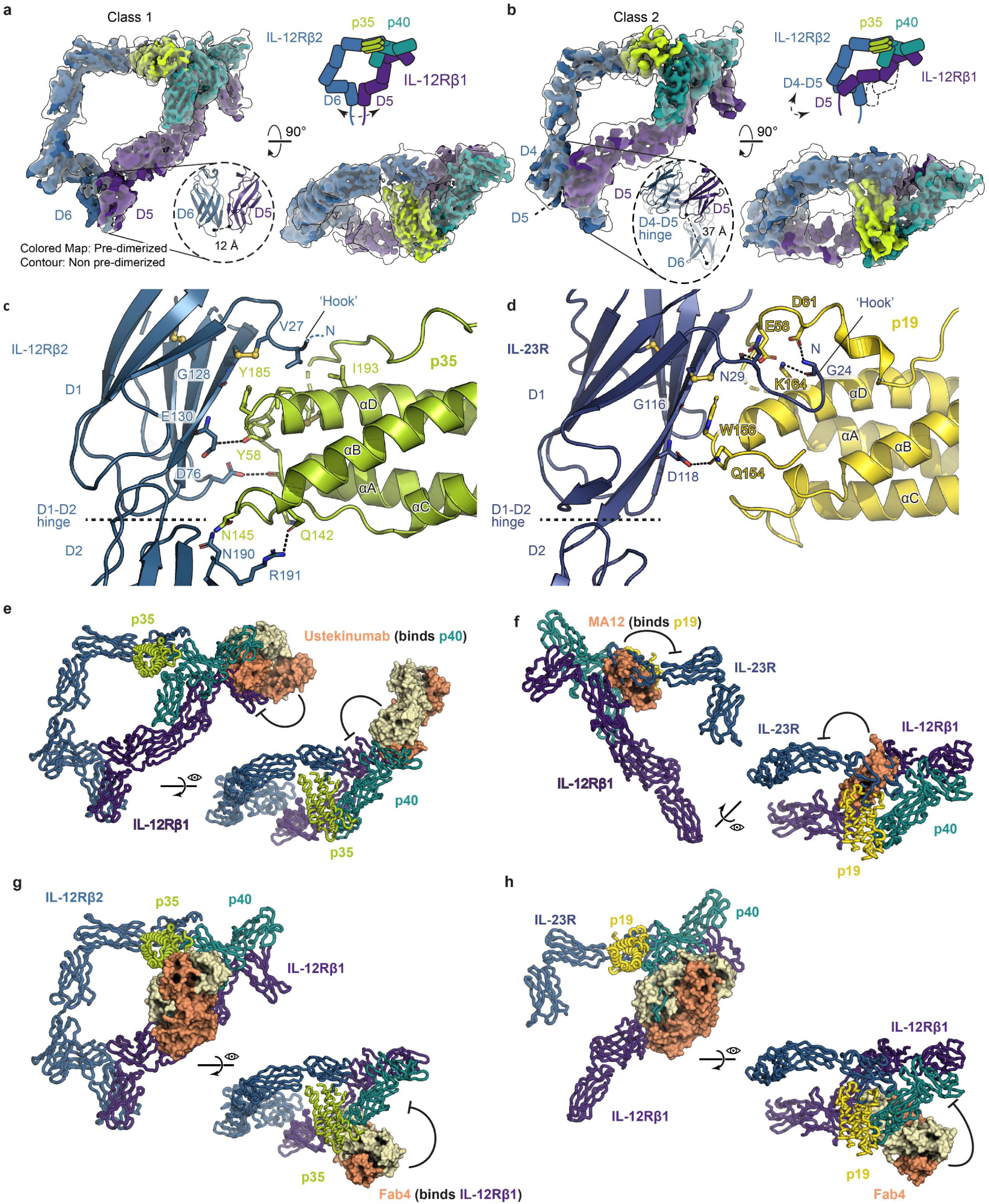
Mechanistic insights into receptor activation by IL-12/IL-23 and antagonism of the common IL-12Rβ1 by Fab4. **a & b**, 3D Class 1 (a) and Class 2 (b) of the pre-dimerized (containing DAPK1_300-319_ & CaM tags) and non pre-dimerized (containing Strep-II tags) extracellular mIL-12:mIL-12Rβ1_D1-D5_:mIL-12Rβ2_D1-D6_ complex. The pre-dimerized Class 1 (a) and Class 2 (b) maps are shown colored in green (mIL-12A/p35), cyan (mIL-12B/p40), purple (mIL-12Rβ1_D1-D5_-DAPK1) and blue (mIL-12Rβ2_D1-D5_-CaM), while the non pre-dimerized Class 1 (a) and Class 2 (b) maps are shown in transparent light grey. Schematic representations of the Class 1 (a) and Class 2 (b) complex assemblies are shown on the top right of panel a and b. **c**, Cartoon representation of the mIL-12A/p35 – IL-12Rβ2 interface. Interacting residues are annotated and shown as sticks, and polar contacts are represented as black dotted lines. mIL-12A/p35 and IL-12Rβ2 are colored green and blue respectively. **d**, Cartoon representation of the hIL-23A/p19 – IL-23R interface. Interacting residues are annotated and shown as sticks, and polar contacts are represented as black dotted lines. hIL-23A/p19 and IL-23R are colored yellow and dark blue respectively. **E-f**, Structural superposition of hIL-12 bound Ustekinumab Fab (pdb 3hmx) and hIL-23 bound alphabody MA12 (pdb 5mj3) (only antagonist shown in surface representation) onto the mIL-12 and hIL-23 ternary complexes (cartoon representation). **G & h** Structural superpostion of hIL-12Rβ1 bound to Fab4 (surface representation) onto the mIL-12 and hIL-23 ternary complexes (cartoon representation).

**Table 1.** Cryo-EM data collection, refinement and validation statistics.

|  | Pre-dimerized<br>mIL-12 complex,<br>Class 1<br>(EMDB-16820)<br>(PDB 8odz) | Pre-dimerized<br>mIL-12 complex,<br>Class 2<br>(EMDB-16821)<br>(PDB 8oe0) | Non pre-dimerized<br>mIL-12 complex,<br>Class 1<br>(EMDB-16822) | Non pre-dimerized<br>mIL-12 complex,<br>Class 2<br>(EMDB-16823) | Pre-dimerized<br>IL-23 complex<br>(EMDB-16824)<br>(PDB 8oe4) |
| --- | --- | --- | --- | --- | --- |
| **Data collection and processing** |  |  |  |  |  |
| Magnification | 60,000 | 60,000 | 60,000 | 60,000 | 60000 |
| Voltage (kV) | 300 | 300 | 300 | 300 | 300 |
| Electron exposure (e<sup>-</sup>/Å<sup>2</sup>) | 61.8 | 61.8 | 61.8 | 61.8 | 61.8 |
| Defocus range (μm) | 0.8 - 3.0 | 0.8 - 3.0 | 0.8 - 3.0 | 0.8 - 3.0 | 0.8-2.0 |
| Pixel size (Å) | 0.755 | 1.51 | 1.51 | 1.51 | 0.9815 |
| Symmetry imposed | C1 | C1 | C1 | C1 | C1 |
| Initial particle images (no.) | 1497836 | 1497836 | 538128 | 538128 | 1454402 |
| Final particle images (no.) | 268498 | 211151 | 66809 | 53211 | 326244 |
| Map resolution (Å) | 3.64 | 4.63 | 5.61 | 6.86 | 3.50 |
| FSC threshold | 0.143 | 0.143 | 0.143 | 0.143 | 0.143 |
| Map resolution range (Å) | 9.0 - 3.0 | 10.0 - 4.0 | 11.0 - 4.5 | 12.0 - 5.5 | 9.0-2.8 |
| **Refinement** |  |  |  |  |  |
| Initial model used (PDB code) | 8cr6 | 8odz | N.A. | N.A. | 5mxa,<br>6wdp,<br>8c7m |
| Model resolution (Å) | 3.90 | 6.50 |  |  | 3.80 |
| FSC threshold |  |  |  |  |  |
| Model resolution range (Å) | ∞ - 3.90 | ∞ - 6.50 |  |  | ∞ - 3.80 |
| Map sharpening B factor (Å<sup>2</sup>) | 106.8 | 259.8 |  |  | 139.6 |
| Model composition |  |  |  |  |  |
| Non-hydrogen atoms | 12,105 | 12,003 |  |  | 9955 |
| Protein residues | 1,515 | 1,503 |  |  | 1249 |
| Ligands | NAG: 7,<br>BMA: 1,<br>MAN: 1 | NAG: 7<br>BMA: 1, |  |  | NAG : 6<br>BMA : 1<br>MAN :1 |
| B factors (Å<sup>2</sup>) |  |  |  |  |  |
| Protein | 91.17 | 317.03 |  |  | 250.33 |
| Ligand | 80.51 | 224.44 |  |  | 163.28 |
| Water | 23.25 | N.A. |  |  | N.A. |
| R.m.s. deviations |  |  |  |  |  |
| Bond lengths (Å) | 0.006 | 0.005 |  |  | 0.008 |
| Bond angles (°) | 1.209 | 0.919 |  |  | 1.128 |
| Validation |  |  |  |  |  |
| MolProbity score | 1.77 | 1.82 |  |  | 1.43 |
| Clashscore | 7.99 | 11.52 |  |  | 4.87 |
| Poor rotamers (%) | 0.00 | 0.30 |  |  | 0.55 |
| CaBLAM outliers (%) | 1.30 | 1.25 |  |  | 1.07 |
| CC (mask/volume) | 0.67/0.66 | 0.70/0.69 |  |  | 0.77/0.76 |
| EMRinger |  |  |  |  |  |
| (sharpened/unsharpened) | 1.26/1.17 | 0.72/0.42 |  |  | 1.55/2.19 |
| Ramachandran plot |  |  |  |  |  |
| Favored (%) | 95.16 | 96.40 |  |  | 96.99 |
| Allowed (%) | 4.84 | 3.60 |  |  | 3.01 |
| Disallowed (%) | 0.00 | 0.00 |  |  | 0.00 |
| Ramachandran Z-score |  |  |  |  |  |
| Whole | -0.99 | -0.85 |  |  | -0.17 |
| Helix | 0.52 | 0.38 |  |  | 0.20 |
| Sheet | 0.19 | 0.27 |  |  | 0.34 |
| Loop | -1.48 | -1.43 |  |  | -0.49 |

A comparison of unbound mIL-12 (X-ray structure of mIL-12, this paper), hIL-12 extracted from an hIL-12:Fab complex (PDB code: 3hmx^12^), and mIL-12 in complex with its two cognate receptors (this paper, cryo-EM Classes 1 and 2) reveals considerable flexibility in D1 of IL-12B (p40), as well as in the tilt of the IL-12A (p35) four-helix bundle around a pivot located at the interface with IL-12B_D2-D3_ (Extended Data Figure 5a). Binding of IL-12 to the D1-D2 hinge of IL-12Rβ1 results in reorientation of D1 of IL-12B to a receptor-bound conformation (Extended Data Figure 5a). Similarly, binding of IL-12 to IL-12Rβ2 fixes the position of the IL-12A α-helices around the IL-12B D2-D3 hinge (Extended Data Figure 5b). For IL-23, however, structural changes in IL-23A (p19) (PDB code: 5mxa^14^) upon receptor binding (this paper and PDB code: 6wdq^15^) are more subtle and local in nature. Whereas the overall position of the four-helix bundle of IL-23A remains the same upon receptor binding, in contrast to IL-12A, its interaction with IL-23R selects for a transition at the N-terminal end of helix αD from an α- to a 3_10_-helix (Extended Data Figure 5c,d).

To date, the interaction interfaces between IL-12 and its two cognate receptors had remained poorly understood, in contrast to IL-23:receptor interactions. The herein presented cryo-EM maps reveal the IL-12:receptor interaction interfaces to ~3 Å resolution (Extended Data Figure 4a,b) and enable head-to-head comparisons between the complexes mediated by IL-12 and IL-23 (Figure 2c,d, Extended Data Figure 6,7,8). Although the mIL-12A:IL-12Rβ2 interaction interface is less extensive than the hIL-23A:IL-23R interface (~700 Å^2^ vs ~900 Å^2^), both interfaces display remarkable commonalities. These include the presence of an N-terminal hook-like structure in domain 1 (D1) of IL-12Rβ2/IL-23R interacting with helix αD of IL-12A/IL-23A, the latter transitioning at the start of the helix to a 3_10_-helix and projecting conserved aromatic residues at its N-terminal end (Tyr185 in mIL-12A and Trp156 in hIL-23A) to stack against the peptide plane of glycine residues in D1 of IL-12Rβ2/IL-23R (Gly128 in mIL-12A and Gly116 in hIL-23A) (Figure 2c,d; Extended Data Figure 6,7,8). Notably, mutation of Tyr185 in mIL-12 and the related Tyr189 in hIL-12 fully abolish binding to IL-12Rβ2^18^. Thus, considering the herein presented structural insights, these positions serve as functional hotspots in IL-12 analogous to the role of Trp157/156 in m/hIL-23^14^. Thus, both IL-12 and IL-23 utilize an aromatic residue on their α-subunit as a hotspot for their interaction with the N-terminal Ig-domain of their high affinity receptors. However, clear differences between the IL-12/IL-23-receptor interfaces are also apparent. The N-terminal hook in IL-12Rβ2_D1_ is much shorter compared to IL-23R and only contacts the tip of α-helix D and stays clear of the loops hanging above it (Figure 2c,d). Furthermore, while IL-23A exclusively interacts with D1 of IL-23R, Gln142 and Asn145 at the tip of helix αB of IL-12A make additional interactions with Asn190 and Arg191 of D2 of IL-12Rβ2, centered around the D1-D2 hinge (Figure 2c,d; Extended Data Figure 6,7,8). These additional interactions appear to compensate for the shorter N-terminal hook of IL-12Rβ2, which only contributes ~110 Å^2^ of interface area, compared to 400 Å^2^ for the corresponding N-terminal hook of IL-23R.

The two cryo-EM 3D classes obtained for mIL-12 in complex with cognate receptors that have been engineered with heterodimerization tags both revealed previously uncharacterized receptor-receptor contacts between the membrane proximal domains of IL-12Rβ1 and IL-12Rβ2 (Figure 2a,b; Extended Data Figure 4a). Remarkably, these 3D classes reveal two distinct contacts between the membrane proximal domains of IL-12Rβ1 and IL-12Rβ2. Whereas Class 1 is characterized by interactions between β-strands βA/βG of IL-12Rβ1 domain 5 and βA/βG of IL-12Rβ2 domain 6, Class 2 features a shift in D4-D5 of IL-12Rβ1 resulting in contacts between IL-12Rβ1 domain 5 and the D4-D5 hinge of IL-12Rβ2 (Figure 2a,b). As our structures carried engineered DAPK1/CaM heterodimerization tags at the C-terminal receptor ends and even though the tags were completely invisible in the cryo-EM maps, it is still possible that the employed tags might have influenced the observed receptor-receptor contacts. To rule out this possibility, we co-expressed mIL-12 with IL-12Rβ1 and IL-12Rβ2 devoid of any dimerization tag in HEK293S cells and characterized their ensuing ternary complex by cryo-EM (Methods). This revealed two 3D classes of the mIL-12 ligand-receptor complex identical to the pre-dimerized complexes, albeit at substantially lower resolutions of 5.6 Å for Class 1 and 6.9 Å for Class 2 (Extended Data Figure 4c). The observed lower resolution of this analysis is presumably due to the increased conformational heterogeneity of the complex, the lower number of particles in the final 3D refinements, and lower number of different particle orientations (Extended Data Figure 4c). The latter are apparent from representative 2D classes after 2D classification (Figure 1c) and the anisotropic character of the refined 3D maps (Extended Data Figure 4c). It is unclear which of the two (or both) observed 3D classes of the extracellular mIL-12 complex might correspond to an active signaling assembly. However, based on the location of membrane proximal domains of IL-12Rβ1 (D5) and IL-12Rβ2 (D6) in the two resulting models, Class 1 positions the C-termini at ~12 Å apart, which is more likely to result in a productive juxta-positioning of transmembrane and cytoplasmic receptor domains when compared to the intermolecular distance of ~37 Å in Class 2 (Figure 2b).

Contrasting the employment of a much longer high affinity receptor IL-12Rβ2 by IL-12, the analogous receptor for IL-23, IL-23R, has shed the three membrane proximal domains during evolution. Membrane proximal receptor-receptor contacts are therefore not possible nor present in the cryo-EM map of the receptor complex mediated by IL-23 (Figure 1d, Extended Data Figure 4b). However, the receptors do adopt similar conformations as in the IL-12 ligand-receptor complex, but with considerable continuous motion as there is little to constrain them beyond the cytokine binding site (Figure 1d). Nevertheless, we envisage that the corroborated observation of two distinct classes of receptor-receptor interactions mediated by binding of IL-12 offers new structural insights into biologically relevant interaction interfaces for therapeutic targeting.

Currently, several antibodies and non-antibody scaffolds binding either IL-12 or IL-23 have been developed with a clear mode of action^12,14, 19–21^ (Figure 2e-f). Intriguingly, a class of antagonistic antibodies against IL-12Rβ1 bind this receptor at an interface that does not overlap with the cytokine binding site (patent WO2012045703A1, Novartis). To gain insights into this antibody’s antagonistic nature, we determined the structure of its antigen-binding portion (Fab4) in complex with domains 3 to 5 (D3-D5) of hIL-12Rβ1 (Methods). The resulting crystal structure at 2.6 Å resolution reveals that Fab4 binds IL-12Rβ1 at the interface of D3 and D4, which is indeed not part of a cytokine- or receptor-receptor interface (Figure 2f,g; Extended Data Figure 9). The paratope of Fab4 is rich in aromatic residues at the periphery, capturing at its center Arg299 of IL-12Rβ1 via hydrogen bonds to the carboxyl groups of Asp105 in the heavy chain and Asp32 in the light chain (Extended Data Figure 9). The Fab4 IL-12Rβ1 interface also traps a few water molecules, of note the water molecule captured between light chain CDR3 and the β4-β5 turn of IL-12Rβ1 D3. When aligning the Fab4-bound structure to the IL-12/IL-23:receptor complexes as determined by cryo-EM, we show that the Fab fragment causes steric hindrance with D3 of IL-12B/p40 in both IL-12 and IL-23. Thus, Fab4 and cytokine binding to IL-12Rβ1 is mutually exclusive (Figure 2g,h).

Collectively, the structures presented in this study provide the long-sought insights into the complete extracellular receptor complexes mediated by pro-inflammatory IL-12 and IL-23 to enable their further mechanistic interrogation and therapeutic targeting.

## Methods

### Cloning and plasmid construction

Cloning was performed by traditional restriction ligation. Q5 DNA-polymerase, T4 ligase, calf intestinal alkaline phosphatase and assorted restriction enzymes were all purchased from NEB. Sequence optimized cDNA encoding cytokines and receptors were purchased from GenScript. Primers and other sequence optimized cDNA were purchased from IDT and TWIST.

All constructs except for the aKappa-VHH were destined for heterologous expression in mammalian HEK293 cells and cloned into the pHLsec vector^22^. The aKappa-VHH was cloned into the pEt15b vector. An overview of all constructs can be found in Supplementary Table 2.

For the Crystalkappa version of the Fab4, light chain residues H215-V221 were exchanged for a shorter fragment encoding QGTTS by PCR^23^.

### Recombinant protein expression

Production of most proteins or protein complexes, except those used in the crystallization of mIL-12, hIL-23 and the mIL-12R:mIL-12 Rβ1_D1-D2_ complex, were performed in suspension-adapted HEK293S *MGAT^−/−^*cells (kindly provided by Prof. N. Callewaert, Unit for Medical Biotechnology, VIB-UGent Center for Medical Biotechnology, Ghent, Belgium). HEK293S *MGAT^−/−^* cells were maintained in a 1:1 ratio of Freestyle (ThermoFisher) and Ex-Cell (Sigma-Aldrich) medium. Transient transfection was performed in Freestyle medium at a density of 3 x 10^6^ cells/mL, with linear polyethylenimine (PEI) 25 kDa (Polysciences) as transfection reagent and using 4.5 µg DNA and 9 µg PEI per mL of transfection volume. Ratio’s of pDNA used in co-transfection were as follows:

- pre-dimerized mIL-12 complex (mIL-12a:mIL-12b-Casp3-AviTag-His_6_:mIL-12Rβ1_D1-D5_-DAPK1: mIL-12Rβ2_D1-D6_-CaM): 4:2:3:4
- non-pre-dimerized mIL-12 complex (mIL-12a:mIL-12b-Casp3-AviTag-His_6_:mIL-12Rβ1_D1-D5_-Strep-II Tag:mIL-12Rβ1_D1-D6_-Strep-II Tag): 4:2:9:4
- pre-dimerized hIL-23 complex (hIL-23_T2A:hIL-12Rβ1_D1-D5_-DAPK1:hIL-23R-CAM): 1:1:1
- hIL-12Rβ1_D3-D5_:Fab4_CrystalKappa_ complex (hIL-12Rβ1_D3-D5_:Fab4 Light_CrystalKappa_:Fab4 Heavy): 3:2:1
- hIL-12Rβ1_D3-D5_:Fab4 complex (hIL-12Rβ1_D3-D5_:Fab4 Light:Fab4 Heavy:aKappa-VHH): 1:1:1
- hIL-23 (hIL-23a:hIL-12b): 1:1

After 5 hours of co-transfection, an equal volume of Ex-Cell medium supplemented with penicillin/streptomycin mix (P/S: 10^6^ units/L penicillin G, 1 g/L streptomycin) was added to the transfected cells, resulting in a density of 1.5 x 10^6^ cells/mL. On day 3 post-transfection, valproic acid and glucose were added to the cells resulting in final concentrations of 2 mM and 27 mM respectively. Human IL-23 was expressed similarly as described above in suspension adapted HEK293S cells.

Production of murine IL-12, mIL-12Rβ1_D1-D2_-CASP3-AviTag-His_6_ and mIL-12b_C197S_-His_6_ in complex with mIL-12Rβ1_D1-D2_-His_6_ was performed in adherently grown HEK293S MGAT^−/−^ cells, seeded in 5-layer multi-flasks (Corning) and grown in DMEM medium supplemented with 10% Fetal Calf Serum (FCS). Transient transfection was performed at a cell confluency of approximately 80%, with branched PEI 25 kDa (Sigma) as transfection reagent and using 50 µg DNA and 75 µg PEI per 175 cm² of transfection surface. For mIL-12, plasmids containing mIL-12a-AviTag-His_6_ and untagged mIL-12b were co-transfected using a plasmid ratio of 1:3, while for mIL-12b_C197S_-His_6_ in complex with mIL-12Rβ1_D1-D2_-His_6_ a plasmid ratio of 1:1 was used. Before adding the DNA-PEI mix, medium was replaced by DMEM without FCS, supplemented with 3.6 mM valproic acid.

aKappa-VHH was expressed in *E. coli* BL21 lysY/Iq (NEB cat n° C3013). The expression plasmid was transformed via heat-shock followed by selection of clones on LB-Agar plates supplemented with 100 µg/mL carbenicillin. An ON grown preculture was used to inoculate the expression culture in LB supplemented with 100 µg/mL carbenicillin and 25 mM sodium phosphate pH 7.2 at 37° C. Expression was induced with 1 mM IPTG and left to express ON at 301 K.

### Recombinant protein purification

#### mIL-12 ligand-receptor complexes for cryo-electron microscopy (cryo-EM)

After 7 days of expression, expression medium was harvested, centrifuged for 5 min at 4000 g to get rid of cellular debris, and filtered using a 0.22 µM bottle-top vacuum filter (Steritop). Clarified and filtered medium containing mIL-12 in complex with either Strep-II tagged mIL-12Rβ1_D1-D5_ and mIL-12Rβ2_D1-D6_ or mIL-12Rβ1_D1-D5_-DAPK1 and mIL-12Rβ2_D1-D6_-CaM, was purified via Immobilized Metal Affinity Chromatography (IMAC) using a prepacked 5 mL cOmplete^TM^ His-Tag purification column (Roche) equilibrated in HEPES-buffered saline buffer (HBS, 25 mM HEPES, pH 7.4, 150 mM NaCl) with or without 5 mM CaCl_2_ for mIL-12 complexes with DAPK1/CaM or Strep-II Tag mIL-12R constructs respectively. After elution using HBS supplemented with 200 mM Imidazole, fractions containing mIL-12 complexes were first desalted using a HiPrep™ 26/10 Desalting column (Sigma-Aldrich) equilibrated in HBS, and subsequently subjected to an overnight Capsase-3 digest (1/100 w/w) at 287 K. The next day, 15 mM Imidazole was added to the Capsase-3 cleaved sample, and an additional IMAC step was performed using a prepacked 5 mL Ni-NTA Fast Flow column (Cytiva) with identical buffers as the previous IMAC step. After washing the column with HBS + 50 mM Imidazole, elution was performed with HBS + 300 mM Imidazole. The flow-through and 50 mM Imidazole wash fractions were combined, concentrated using a 15 kDa Amicon® Ultra-15 Centrifugal Filter Unit (Merck) and injected onto a Superdex 200 HiLoad 16/600 column (GE Healthcare) equilibrated in HBS supplemented with or without 5 mM CaCl_2_ for mIL-12 complexes with DAPK1_302-330_/CaM or Strep-II Tag mIL-12R constructs respectively. Complexes containing mIL-12 and Strep-II tagged mIL-12R constructs were subjected to an additional polishing Size Exclusion Chromatography (SEC) step after pooling the left fractions of the peak after the first SEC run, using a Superdex 200 Increase 10/300 GL column equilibrated in HBS. After the final SEC, peak fractions were aliquoted and frozen in liquid nitrogen for further use.

#### hIL-23 ligand-receptor complex for cryo-EM

Recombinant protein was captured from the clarified and filtered conditioned medium by IMAC (equilibration buffer: HBS supplemented with 5 mM CaCl_2_, elution buffer: HBS supplemented with 5 mM CaCl_2_ and 200 mM imidazole). Protein complexes were captured via the only His-tag present on the p19 subunit of the cytokine. To remove potential soluble aggregates the IMAC eluate was separated by SEC on a Superose6 increase 10/300 GL column (Cytiva) (running buffer: HBS supplemented with 5 mM CaCl_2_). The seemingly monomeric complexes were pooled and digested overnight with Caspase-3 (1/100 w/w) to remove the His-tag on the cytokine. A second IMAC was performed to capture the undigested cytokine, recombinant Caspase-3 and potential contaminants interacting with the IMAC resin. Finally, a polishing SEC was performed on a Superdex200 Increase 10/300 GL column (running buffer: HBS supplemented with 5 mM CaCl_2_). The top fractions were aliquoted and frozen in liquid nitrogen for storage before further use.

#### mIL-12 and mIL-12b_C197S_:mIL-12Rβ1_D1-D2_ complex for X-ray crystallography

After 4 days of expression, medium was harvested, centrifuged to get rid of cellular debris, and filtered prior to loading onto a prepacked 5 mL cOmplete^TM^ His-Tag purification column (Roche) equilibrated in HBS. After elution using HBS + 300 mM Imidazole, fractions containing mIL-12b_C197S_-His_6_ in complex with mIL-12Rβ1_D1-D2_-His_6_ were concentrated and injected onto a Superdex 200 Increase 10/300 GL column equilibrated in HBS. Purified mIL-12b_C197S_-His_6_ in complex with mIL-12Rβ1_D1-D2_-His_6_ was subjected to an overnight Caspase-3 and EndoH (New England Biolabs) digest (1/100 w/w) at 287 K, and subsequently purified using a second and final SEC step using a Superdex 200 Increase 10/300 GL column equilibrated in HBS. Purified, deglycosylated mIL-12b_C197S_:mIL-12Rβ1_D1-D2_ complex was concentrated to approximately 7.4 mg/mL, aliquoted, and flash-cooled in liquid nitrogen for further use.

For the expression of mIL-12, the same protocol was used as described for the mIL-12b_C197S_:mIL-12Rβ1_D1-D2_ complex, except for the addition of Capsase-3 (1/100 w/w) during overnight EndoH treatment, and the inclusion of a negative IMAC step using a HisTrap IMAC HP column (GE Healthcare) before performing the final SEC run. Purified, deglycosylated mIL-12 was concentrated to approximately 6.4 mg/mL, aliquoted, and flash-cooled in liquid nitrogen for further use.

#### hIL-12Rβ1_D3-D5_:Fab4_CrystalKappa_ and hIL-12Rβ1_D3-D5_:Fab4:anti-Kappa-VHH complexes for X-ray crystallography

For the VHH purification, cells were harvested by centrifugation, lysed by 3 passages in a high pressure homogenizer (Emulsiflex) and clarified by centrifugation and filtration. aKappa-VHH was captured by IMAC followed by SEC on a HiLoad Superdex 75 16/600. The His tag was removed by overnight digest with Caspase-3 (1/100 w/w) at RT. Following a second IMAC to capture undigested VHH, recombinant enzyme and contaminants, the tagless VHH was finally polished by a second SEC separation and frozen in liquid nitrogen for storage before further use.

For the coexpressed hIL-12Rβ1_D3-D5_:Fab4_CrystalKappa_ and hIL-12Rβ1_D3-D5_:Fab4 complexes, the recombinant protein was purified from the clarified and filtered conditioned media by IMAC followed by SEC. Only the SEC fractions containing an apparent monomeric complex were propagated further. After trimming of the N-linked glycans by ON digest with EndoH (1/100 w/w) together with cleavage of the His tags with Caspase-3 a second IMAC was performed to capture undigested VHH, recombinant enzyme and contaminants. A final polishing step was performed was performed on a Superdex200 Increase 10/300 GL column (running buffer: HBS). For the hIL-12Rβ1_D3-D5_:Fab4:anti-Kappa-VHH complex, a molar excess of previously purified VHH was added to the tagless and deglycosylated receptor Fab complex prior to injection on SEC.

#### hIL-23 for X-ray crystallography

hIL-23 was purified from the clarified and filtered conditioned medium by IMAC followed by SEC on a Superdex200 Increase 10/300 GL column (running buffer: HBS). The His tag was not removed from the cytokine prior to crystallization. The final yield was around 100 mg/L expression medium.

#### mIL-12 receptor constructs for Bio-Layer Interferometry (BLI)

Purification of mIL-12Rβ1_D1-D2_-CASP3-AviTag-His_6_ was performed using the same protocol as described for mIL-12, but without adding EndoH during the overnight Caspase-3 digest. For purification of mIL-12Rβ2_D1-D6_-CASP3-AviTag-His_6_, centrifuged and filtered expression medium was loaded onto a prepacked 5 mL cOmplete^TM^ His-Tag purification column (Roche) equilibrated in HBS. After elution using HBS + 300 mM Imidazole, fractions containing mIL-12Rβ2_D1-D6_-CASP3-AviTag-His_6_ were desalted using a HiPrep™ 26/10 Desalting column (Sigma-Aldrich) equilibrated in HBS, concentrated, and subsequently injected onto a Superdex 200 HiLoad 16/600 column (GE Healthcare) equilibrated in HBS.

### Cryo-electron microscopy (cryo-EM)

For each complex, a 4 µl sample of purified complex was applied to glow-discharged R2/1 300 mesh holey carbon copper grids (Quantifoil Micro Tools GmbH), using a conc. of 0.13 and 0.09 mg/ml for the pre-dimerized and non pre-dimerized mIL-12 complex respectively, and using a conc. of 0.40 mg/ml for the pre dimerized hIL-23 complex. Grids were blotted and plunge frozen in liquid ethane using a Lecia EM GP2 Plunge Freezer operated at 95% humidity, using a blotting time of 5 and 4 s. ml for the pre-dimerized and non pre-dimerized mIL-12 complex, and a blotting time of 4.5 s for the pre dimerized hIL-23 complex. All cryo-EM datasets were recorded at the VIB-VUB facility for Biological Electron Cryogenic Microscopy (BECM, Brussels, Belgium), on a 300 kV CryoARM300 microscope (JEOL) equipped with an Omega filter (JEOL) and K3 direct electron detector (Gatan) operated in correlated double-sampling (CDS) mode. 60-frame movies were collected respectively, with a total exposure of 3.37 s, a total dose of 61.8 e^−^/Å^2^, an energy filter slit width of 20 eV, and at a magnification of 60,000 ×, corresponding to a pixel size of 0.755 Å/pixel at the specimen level. For the pre-dimerized and non pre-dimerized mIL-12 complex a total of 8145 and 7174 movies where collected respectively, while for the pre-dimerized hIL-23 complex 6660 movies were collected.

### Cryo-EM Image Processing

#### mIL-12 ligand-receptor complex

Processing of recorded movies (non pre-dimerized dataset: 7174, pre-dimerized dataset: 8145) was performed in cryoSPARC v3.3.2^24,25^. Movies were motion corrected and dose weighted using Patch motion correction, and CTF estimation on the aligned and dose weighted micrographs was performed using Patch CTF estimation.

For the pre-dimerized dataset, particle picking was initially performed using Blob picker. Inspected particle picks were extracted using a box size of 440 pixels, downsampled to a box size of 220 pixels, corresponding to a pixel size of 1.51 Å/pixel (2x binned). The extracted particle stack was first cleaned using several rounds of iterative 2D classification and 2D class selection. The cleaned particle stack was used as an input for *Ab Initio* model generation using 3 classes as input, followed by heterogeneous refinement of the 3 *Ab Initio* volumes. Since one resulting class clearly corresponded to a partial mIL-12 complex lacking one of the two receptor legs, only two of the three classes were further refined using Non-Uniform (NU) refinement. The first class after NU 3D refinement corresponded to a complete mIL-12 ligand-receptor complex and was used as a template for Template picking. The resulting particle stack was cleaned using iterative 2D classification and 2D class selection and was used for heterogeneous refinement using 3 *Ab Initio* volumes as input, followed by 3D classification without alignment. Two of the three resulting 3D classes corresponded to complete mIL-12 ligand-receptor complexes, and were used for a second cycle of 3D classification without alignment. The two resulting 3D classes were used for NU 3D refinement, resulting in two maps with resolutions of 3.7 Å and 4.6 Å based on the 0.143 gold-standard Fourier shell correlation (FSC) criterion^26^. Particles corresponding to the first map were re-extracted in a box size of 440 pixels corresponding to a pixel size of 0.755 Å/pixel, and used for a final NU 3D refinement, resulting in a final map with a resolution of 3.6 Å based on the 0.143 gold-standard Fourier shell correlation (FSC) criterion. Final maps were post-processed using DeepEMhancer^27^ for the purpose of model building and visualization.

For the non pre-dimerized dataset, particles were picked using crYOLO^28^. The crYOLO picked particle stack was imported in cryoSPARC v3.3.2 and extracted using a box size of 440 pixels, downsampled to a box size of 220 pixels, corresponding to a pixel size of 1.51 Å/pixel (2x binned). The extracted particle stack was cleaned using several rounds of iterative 2D classification and 2D class selection, and the cleaned particle stack was used as an input for *Ab Initio* model generation using 2 classes as input. The obtained two *Ab Initio* models were used as input for heterogeneous refinement (2 classes) followed by a final NU 3D refinement of each class, resulting in two maps with resolutions of 5.6 Å and 6.9 Å based on the 0.143 gold-standard Fourier shell correlation (FSC) criterion.

#### hIL-23 ligand-receptor complex

A comprehensive overview of the data processing can be found in Extended Figure 3.

cryoSPARC v3.3.2 was used to process the 6660 recorded hIL-23 complex movies with a pixel size of 0.755 Å/pixel. Movies were motion corrected and dose weighted using Patch motion correction, and CTF estimation on the aligned and dose weighted micrographs was performed using Patch CTF estimation.

Particle picking was sequentially performed by blob picking, template picking and finally TOPAZ^29^ picking using a model trained on already identified IL-23 complex particles.

Several rounds of 2D classification and heterogenous refinement steps were performed to select a coherent subset of complex particles. This cleaned particle stack was finally subjected to 3D variability analysis allowing for 2 modes. The first mode displayed continuous variability at the level of the receptor legs and was used to select a more homogenous final subset of 326244 particles.

Most of the analysis and the final map was reconstructed with particles were extracted from the micrographs using a box size of 416 pixels and downsampled to 320 pixels corresponding to a pixel size of 0.9815 Å/pixel. The final NU 3D refinement resulted in a map with resolutions of 3.50 Å based on the 0.143 gold-standard Fourier shell correlation (FSC) criterion.

### Cryo-EM Model Building and Refinement

#### mIL-12 ligand-receptor complex

Initial models for mIL-12 were taken from the mIL-12 crystal structure determined herein, and from Alphafold2^17^ structural predictions of mIL-12Rβ1 (https://alphafold.ebi.ac.uk/entry/Q60837) and mIL-12Rβ2 https://alphafold.ebi.ac.uk/entry/P97378). Alphafold2 models were first trimmed to exclude parts with low per-residue confidence score (pLDDT<50). Next, the crystal structure of mIL-12 and trimmed Alphafold2 models of mIL-12Rβ1 and mIL-12Rβ2 were rigid-body fitted in the deepEMhancer sharpened map of pre-dimerized mIL-12 complex Class 1, using USCF Chimera^30^, after which the rigid-body fitted model was subjected to automatic molecular dynamics flexible fitting using NAMDINATOR^31^. The resulting flexibly fitted mIL-12 complex was then used for several rounds of manual building in Coot^32^ followed by cycles of real-space refinement in Phenix 1.19.2-4158^33^ using global minimization, local grid search, atomic displacement parameter (ADP) refinement, secondary structure restraints and Ramachandran restraints. A final refinement was performed using global minimization, local grid search, ADP refinement, secondary structure and Ramachandran restraints, and using a non-bonded weight parameter of 300

Afterwards, the refined structural model of pre-dimerized mIL-12 complex Class 1 was fitted in the deepEMhanced map of pre-dimerized mIL-12 complex Class 2 using ProSMART^34^ restraints in Coot. The resulting fitted structure was used as a starting point for a few rounds of manual building in Coot followed by real-space refinement in Phenix 1.19.2-4158 using global minimization, local grid search, ADP refinement, secondary structure, reference model and Ramachandran restraints. A final refinement was performed using global minimization, local grid search, ADP refinement, secondary structure, reference model and Ramachandran restraints, and using a non-bonded weight parameter of 300. A summary of cryo-EM data collection, refinement and validation statistics can be found in Table 1.

#### hIL-23 ligand-receptor complex

Initial models for the hIL-23 complex were assembled from previously published crystal structures (pdb 5mzv, 6wdp), the crystal structure of the partial hIL-12Rβ1_D3-D5_ presented herein and Alphafold2 structural prediction. These models were rigid body fit into the deepEMhacer sharpened map using USCF ChimeraX. Further restrained flexible fitting was performed in Coot utilizing ProSMART restraints. Iterative model building and refinement was performed in ISOLDE, Coot and Phenix using a non-bonded weight parameter of 200. A summary of cryo-EM data collection, refinement and validation statistics can be found in Table 1.

### X-Ray Crystallography

All crystallization experiments were set up using a Mosquito crystallization robot (SPT Labtech) with sitting drop vapor diffusion geometry in Swissci triple-drop plates.

#### mIL-12

EndoH treated mIL-12 was concentrated to 6.35 mg/mL prior to setting up crystallization trials using 300 nL drops (100 nL protein + 200 nL reservoir solution) at 293 K. Crystals were obtained in a crystallization condition containing 0.2 M TMAO, 0.1 M Tris pH 8.5 and 20% w/v/ PEG 2000 MME (Molecular Dimensions JCSG-plus^TM^, condition G4) supplemented with 20 mM BaCl_2_ (Hampton Research Additive Screen HT condition A1), cryoprotected using mother liquor supplemented with 15% PEG400, and subsequently flash-cooled in liquid nitrogen. Diffraction data were collected at the Proxima2A beamline (SOLEIL synchrotron, Gif-sur-Yvette, France) at 100K, with a beam wavelength of 0.980 Å and size of 50 x 50 µm. Obtained diffraction data were processed using the XDS package^35^.

#### hIL-23

Purified hIL-23 was concentrated to 13 mg/mL prior to crystallization trials. Two hits were identified in the Morpheus2 screen (Molecular Dimensions) containing 2 mM lanthanide mix at pH 7.5 with different precipitants in drops set up with a ration of 1:2 protein to mother liquor. The hit in condition E7 (2 mM lanthanides, 0.1M BES/TEA, 10% PEG8000, 20% 1,5-pentanediol) could be vitrified from the mother liquor in liquid nitrogen prior to data collection. Diffraction data were collected at the P14 beamline (PETRA III, Hamburg, Germany) at 100K.

#### mIL-12B_C197S_:mIL-12Rβ1_D1-D2_ complex

Purified mIL-12B_C197S_ in complex with mIL-12Rβ1_D1-D2_ was concentrated to 7.4 mg/mL before setting up crystallization trials using 200 nL drops (100 nL protein + 100 nL reservoir solution) at 293 K. Crystals were obtained in a crystallization condition containing 20 % PEG Smear Medium (12.5%w/v PEG 3350, 12.5%w/v PEG 4000, 12.5%w/v PEG 2000, 12.5%w/v PEG 5000 MME), 0.1 M MgCl_2_ hexahydrate, 0.1 M KCl and 0.1 M PIPES pH 7.0 (BCS^TM^, condition E4), cryoprotected using mother liquor supplemented with 15% PEG400, and subsequently flash-cooled in liquid nitrogen. Diffraction data were collected at the P13 beamline (PETRA III, Hamburg, Germany) at 100K, with a beam wavelength of 0.98 Å. Obtained diffraction data were processed using the XDS package^35^.

#### hIL-12Rβ1_D3-D5_:Fab4_CrystalKappa_ and hIL-12Rβ1_D3-D5_:Fab4:anti-Kappa-VHH complexes

Attempts at crystallizing hIL-12Rβ1, complexes and fragments thereof mostly failed to yield crystals, or led to poorly diffracting crystals. Purified hIL-12Rβ1_D3-D5_ in complex with wild-type Fab4 also failed to yield crystals. Upon the inclusion of the aKappa-VHH, crystallization trials performed at 287 K with 6.6 mg/ml protein yielded crystals in BCS screen (Molecular dimensions) condition D9 and optimized to 17.27% Tacsimate pH 6.34,10% Ethylene Glycol, 10% PEG Smear Broad. Crystal were cryoprotected in ML supplemented with 17% Ethylene Glycol prior to vitrification in liquid nitrogen. Diffraction data were collected at the P14 beamline with a beam wavelength of 0.92 Å at 100K.

To obtain a qualitative search model to solve the hIL-12Rβ1_D3-D5_:Fab4:aKappa-VHH complex crystal structure, the structure of the Fab4:aKappa-VHH complex was experimentally determined. Purified Fab4:aKappa-VHH was concentrated to 6 mg/ml and crystallization trials set up leading to a hit in Crystal Screen (Hampton research) condition C7. After optimization (180 mM (NH4)_2_SO4, 27% PEG4000), crystals were cryoprotected in ML supplemented with 17% ZW221 (40% v/v DMSO, 40% v/v Ethylene Glycol, 20% v/v Glycerol)^36^. The crystals belong to space group P1 but have an extensive (ca 40%) twin fraction according to the pseudo-merohedral twin operator -h,k,-l. Diffraction data to 2.2Å was used to model the search model used to phase the hIL-12Rβ1_D3-D5_:Fab4:aKappaVHH via molecular replacement. Due to the problematic nature of refining extensively twinned datasets, it was not deposited.

A second crystal of hIL-12Rβ1_D3-D5_ in complex with Fab4 was obtained by utilizing the Crystal Kappa form of the Fab. This Fab mutein mimics the rabbit Kappa light chain which allows for homotypical crystal contacts by beta-sheet complementation. The 12Rβ1_D3-D5_:Fab4_X-talKappa_ protein complex was concentrated to hIL-7 mg/ml. Crystals were identified in Morpheus2 screen condition A3 and were optimized to 90 mM LiNaKSO4, 100 mM Citrate BisTRIS propane pH 5.4, 8.57% PEG 8000, 17.14% 1,5 Pentanediol. Crystals were vitrified from the mother liquor in liquid nitrogen prior to data collection. Diffraction data were collected at the P14 beamline with a beam wavelength of 0.98 Å at 100K.

### X-Ray crystallographic Model Building and Refinement

An overview of all X-Ray data collection and refinement statistics can be found in Supplementary Table 1.

#### mIL-12

Diffraction data for mIL-12 were processed in space group *P* 2_1_ 2_1_ 2_1_ (a=59.11 Å, b=70.77 Å, c=127.37 Å, α=β=γ=90°) to a resolution of 2.9 Å. Structure determination was performed by maximum-likelihood molecular replacement (MR) in Phaser^37^ using the crystal structure of human IL-12 extracted from the hIL-12:Ustekinumab complex (PDB code 3HMX^12^) as a search model. The resulting MR solution was initially refined in Buster^38^ followed by several cycles of manual building in Coot^32^. Final refinements were performed in Phenix 1.19.2-4158^32^ using positional and real-space refinement, individual isotropic ADP refinement combined with TLS (Translation-Liberation-Screw-rotation model) groups, and optimized X-ray/stereochemistry and X-ray/ADP weights. The final model had following Ramachandran statistics: 96.4 % favored, 3.6 % allowed, 0 % outliers.

#### hIL-23

Diffraction data for hIL-23 were processed in space group P 2_1_ (a= 67.75 Å, b= 59.84 Å, c= 69.99 Å,, α=γ=90°, β= 98.06°) to a resolution of 2.0 Å. Initial phases were recovered by maximum-likelihood based molecular replacement in Phaser using a previously determined hIL-23 structure as search model (pdb 5mxa). Model building and refinement was performed iteratively in Coot and Phenix.refine. Extra distance restraints of 2.4 Å between the lanthanide and its interacting partners were imposed. Individual isotropic B-factors were refined together with domain level TLS parameters. The final model had the following Ramachandran statistics: 96.46% favored,3.54% allowed and 0 outliers.

#### mIL-12B_C197S_:mIL-12Rβ1_D1-D2_ complex

Diffraction data for mIL-12B_C197S_ in complex with mIL-12Rβ1_D1-D2_ were processed in space group *P* 1 2_1_ 1 (a=29.22 Å, b=165.09 Å, c=51.87 Å, α=γ=90°, β=90.44°) to a resolution of 2.2 Å. Structure determination was performed by maximum-likelihood MR in Phaser^37^ using the crystal structure of mIL-12B (p40) extracted from a murine p80 homodimer (PDB code 6sff^39^) as a search model. After refinement of the resulting MR solution in in Phenix 1.19.2-4158^33^, additional density appeared for mIL-12Rβ1_D1_, allowing manual building in Coot^32^ and completion of the complex. Western Blot analysis of washed and dissolved mIL-12B_C197S_:mIL-12Rβ1_D1-D2_ complex crystals demonstrate proteolysis starting from the C-terminal His_6_ tag of mIL-12Rβ1_D1-D2_ in the crystallization drops. As a result, density for domain 2 mIL-12Rβ1 is absent in the electron density map. A final refinement was performed in Phenix using positional and real-space refinement, individual isotropic ADP refinement combined with TLS, and optimized X-ray/stereochemistry and X-ray/ADP weights. The final model had following Ramachandran statistics: 96.2 % favored, 3.8 % allowed, 0 % outliers.

#### hIL-12Rβ1_D3-D5_:Fab4_CrystalKappa_ and hIL-12Rβ1_D3-D5_:Fab4:anti-Kappa-VHH complexes

Diffraction data for hIL-12Rβ1_D3-D5_ in complex with Fab4_CrystalKappa_ were processed in space group *P* 2_1_ 2_1_ 2_1_ (a=73.66 Å, b=140.18 Å, c=219.36 Å, α= β =γ=90°) to a resolution of 2.56 Å. Initial phases were recovered by maximum-likelihood based molecular replacement in Phaser using a previously determined Fab4 structure and the AlphaFold2 model for the receptor fragment. The asymmetric unit contains 2 copies of the complex. Model building and refinement was performed iteratively in Coot and Phenix.refine. One copy of the Fab4 has a much lower overall B-factor compared to the other copy, this is thanks to the CrystalKappa mutations allowing for crystal packing interactions to form and limit the movement of the Fab constant domains. Individual isotropic B-factors were refined together with domain level TLS parameters. The final model had following Ramachandran statistics: 97.11 % favored, 2.89 % allowed, 0.00 % outliers.

Diffraction data for hIL-12Rβ1_D3-D5_ in complex with Fab and a kappa light chain recognizing VHH were processed in space group *P* 4_1_ 3 2 (a=b=c=231.40 Å, α= β =γ=90°) to a resolution of 4.40 Å.. Initial phases were recovered by maximum-likelihood based molecular replacement in Phaser using a previously determined Fab4 structure and a published structure of a Fab in complex with the kappa light chain recognizing VHH (pdb 6ana). The mainchain of the receptor fragment could be iteratively traced in coot with intermittent refinement. Interpretable electron density maps could be obtained by refinement in Buster. With the release of the public AlphaFold2 model database model building could be finalized. Residue level group B-factors were refined together with domain level TLS parameters. The final model had following Ramachandran statistics: 95.38 % favored, 4.5 % allowed, 0.12 % outliers.

### Bio-Layer Interferrometry (BLI)

Ligands for BLI studies were prepared using *In Vitro* biotinylation using recombinantly produced BirA enzyme. BLI studies were performed using biotinylated mIL-12 (consisting of mIL-12a-AviTag-His_6_ and untagged mIL-12b) and mIL-12Rβ2_D1-D6_-Casp3-AviTag-His_6_ as the ligands, and mIL-12Rβ1_D1-D2_ and mIL-12 as the respective analytes. Experiments using biotinylated mIL-12 were performed in HBS kinetics buffer (20 mM HEPES pH 7.5, 150 mM NaCl, 0.1% w/v BSA and 0.02% v/v Tween-20), using an Octet RED96 instrument (Sartorius) operated at 298 K. For experiments including biotinylated mIL-12Rβ2_D1-D6_, HBS kinetics buffer was used containing 0.2% w/v BSA and 0.04% v/v Tween-20. Streptavidin-coated biosensors (Sartorius) were functionalized with either biotinylated mIL-12 or biotinylated mIL-12Rβ2_D1-D6_, using loading times of 600 s and 480 s respectively, to reach a signal of ± 1 nm. Next, functionalized sensors were quenched with 10 μg mL^−1^ D-biotin, and transferred to wells containing 5 gradually increasing concentrations of analyte. Buffer subtraction was performed using functionalized biosensors dipped in running buffer. Nonspecific binding was monitored during the experiments using non-functionalized biosensors dipped in the highest ligand concentration as well as running buffer. All data were fitted using the FortéBio (Sartorius) Data Analysis 9.0 software, utilizing a 1:1 interaction model. For all binding experiments three technical replicates were performed, and presented *K*_D_, *k*_d_ and *k*_a_ values are calculated as the averages of these triplicate experiments.

### IL-23 reporter cellular assay

HEK-Blue™ IL-23 Cells (Invivogen, cat° hkb-il23) were maintained in growth medium (Dulbecco’s Modified Eagle Medium, 4.5 g/l glucose, 2 mM L-glutamine and 10% (v/v) heat-inactivated fetal bovine serum) supplemented with 1X HEK-Blue™ Selection antibiotics in a humid chamber with 5 % CO2 in the atmosphere at 310 K. Cells were passaged by trypsin-EDTA digest, at 80% confluency.

For the reporter assay, on the day prior to the cytokine challenge, cells were washed with PBS and detached from the flasks using a cell-scraper. Cells were subsequently seeded in a 96-well plate in a total volume of 180 µL/well at a final density of 2.8×10^5^ cells/mL in growth medium and incubated for 24h. On the day of the cytokine challenge, cells were challenged with a 2.5-fold dilution series of hIL-23 in the presence and absence of 50 nM Fab4 and incubated for a further 24h. The next day, 20 µL of the conditioned medium was added to 180 µL of QUANTI-Blue™ Solution and allowed to incubate for 15 minutes at 310 K. The assay was read out using a spectrophotometer (Bio-Rad) at 655 nm. The concentration of FAb4 used was determined by prior optimization experiments. The assay was performed in triplicate, each with technical triplicates. The readout were globally fit to the Hill equation allowing for one global EC_50_ and Hill coefficient as well as individual R_0_ and R_max_ per biological replicate. The 95% confidence interval was estimated by bootstrap analysis with 1000 replicates. Fitting was performed utilizing the SciPy Python package.

## Data Availability

Cryo-EM maps and accompanying structural models were deposited in the EMDB/PDB with following accession codes: EMD-16820/8odz (pre-dimerized mIL-12 cytokine-receptor complex, Class 1), EMD-16821/8oe0 (pre-dimerized mIL-12 cytokine-receptor complex, Class 2), EMD-16824/8oe4 (pre-dimerized hIL-23 cytokine-receptor complex). Cryo-EM maps of the non pre-dimerized mIL-12 cytokine-receptor complex (Class 1 & 2) were deposited in the EMDB with codes EMD-16822 and EMD-16823 respectively. Crystallographic coordinates and structure factors were deposited to the PDB with following accession codes: 8cr6 (mIL-12), 8cr5 (mIL-12B_C197S_:mIL-12Rβ1_D1-D2_ complex), 8cr8 (hIL-23), 8c7m (hIL-12Rβ1_D3-D5_:Fab4_CrystalKappa_ complex), 8odx (hIL-12Rβ1_D3-D5_:Fab4:anti-Kappa-VHH complex).

## Acknowledgements

We thank the staff of beamlines Proxima2A (SOLEIL synchrotron, Gif-sur-Yvette, France) and P14 (PETRA III, Hamburg, Germany) for beamtime allocation and technical support. We thank Marcus Fislage at the VIB-VUB Facility for Bio Electron Cryogenic Microscopy (BECM) for assistance in data collection, technical support and infrastructural access. Y.B. was a post-doctoral research fellow supported by the Research Foundation Flanders (FWO grant n° 12S0519N). S.N.S. acknowledges research support from the FWO (grant n° G0B4918N) and the Flanders Institute for Biotechnology (VIB).

## Author information

### Authors and Affiliations

Unit for Structural Biology, Department of Biochemistry and Microbiology, Ghent University, Ghent, Belgium

Yehudi Bloch, Jan Felix, Romain Merceron,Mathias Provost, Royan Alipour Symakani, Robin de Backer, Elisabeth Lambert, Savvas N. Savvides

Unit for Structural Biology, VIB-UGent Center for Inflammation Research, Ghent, Belgium

Yehudi Bloch, Jan Felix, Romain Merceron,Mathias Provost, Royan Alipour Symakani, Robin de Backer, Elisabeth Lambert, Savvas N. Savvides

Eurofins DiscoverX Products France, Celle-Lévescault, France

Romain Merceron

VIB Center for Medical Biotechnology, Ghent, Belgium

Royan Alipour Symakani

Solvias, Kaiseraugst, Switzerland

Elisabeth Lambert

Hamburg Unit c/o DESY, European Molecular Biology Laboratory, Hamburg, Germany

Yehudi Bloch

### Contributions

Y.B., J.F. and R.M. designed and performed recombinant protein production with contributions from M.P., R.A.S. and E.L. J.F. and Y.B. performed cryo-electron microscopy grid preparation, data collection, processing, model building and refinement. Y.B, and R.M performed X-ray crystallographic data collection. Y.B., J.F. and R.M. performed X-ray crystallographic data processing, model building and refinement. J.F. and R.A.S performed BLI binding studies. M.P and Y.B. performed IL-23 reporter cellular assays. Y.B., J.F. and S.N.S. analyzed data with contributions from R.D.B. J.F., Y.B. and S.N.S. wrote the manuscript with contributions from all authors. S.N.S. conceived and supervised the project.

### Corresponding authors

Correspondence to Jan Felix or Savvas N. Savvides.

---

**Extended Data Figure 1:**
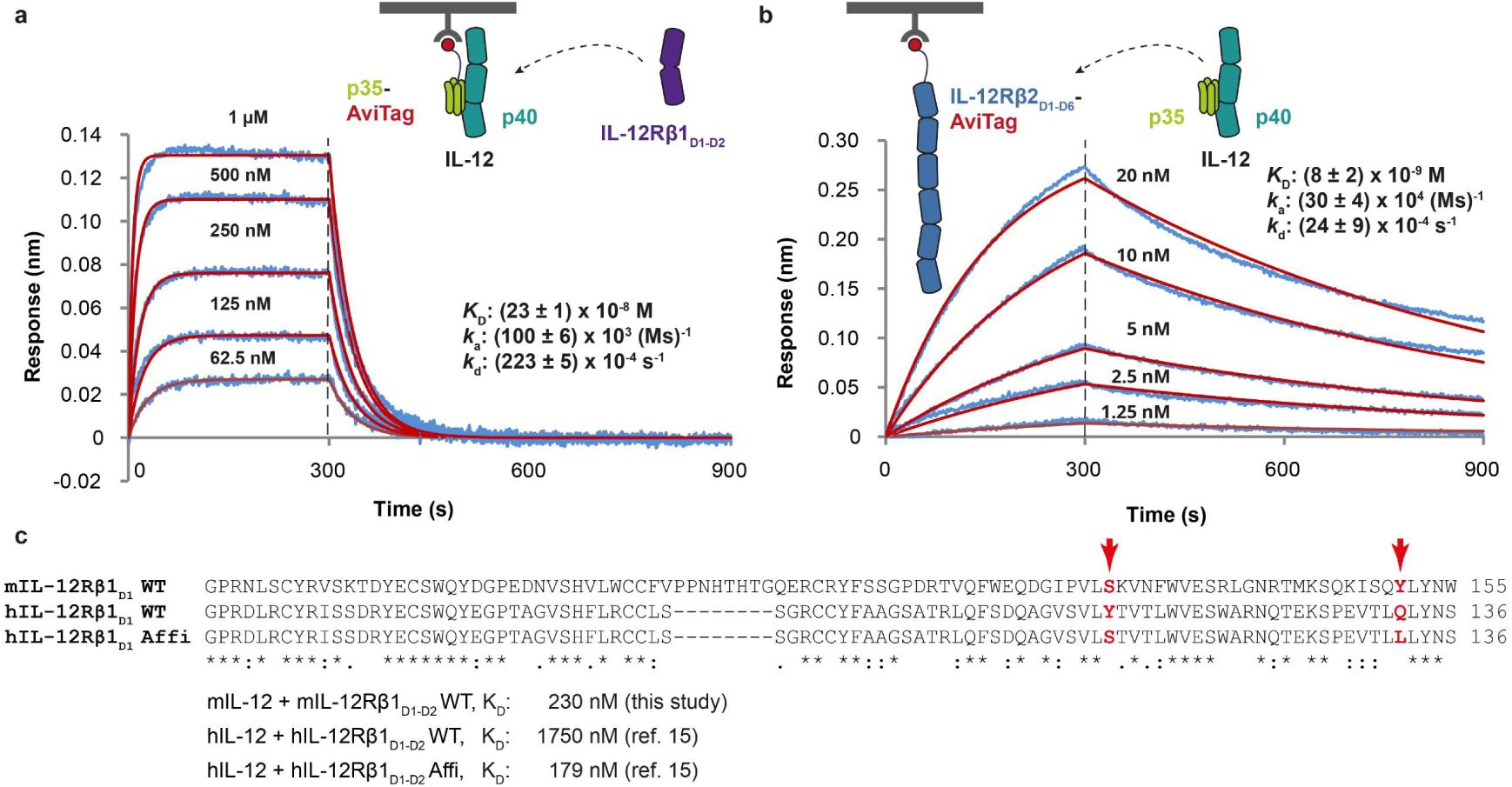
Characterization of the interaction between murine IL-12 and its cognate receptors via Bio-Layer Interferometry (BLI). **a & b**, BLI measurements of the interaction between biotinylated mIL-12, couped on streptavidin (SA) biosensors, and mIL-12Rβ1_D1-D2_ (a) or biotinylated mIL-12Rβ2_D1-D6_ and mIL-12 (b). Schematic representations of the interactions are shown above each set of measurements. Measured response curves are shown in blue, and fitted curves (using a 1:1 binding model) are shown in red. Used concentrations of analyte are annotated above each individual response curve. *K*_D_ = Dissociation constant. All BLI experiments were performed in triplicate (see Source Data file), and one representative experiment is shown. Displayed *K*_D_ values are the calculated average of triplicate experiments. **c,** Multiple sequence alignment of domain 1 (D1) of wild-type hIL-12Rβ1, affinity maturated hIL-12Rβ1^15^ and wild-type mIL-12Rβ1. The location of mutated residues in affinity maturated hIL-12Rβ1 is annotated with a red arrow. Sequence alignment was performed using Clustal Omega^40^.

**Extended Data Figure 2:**
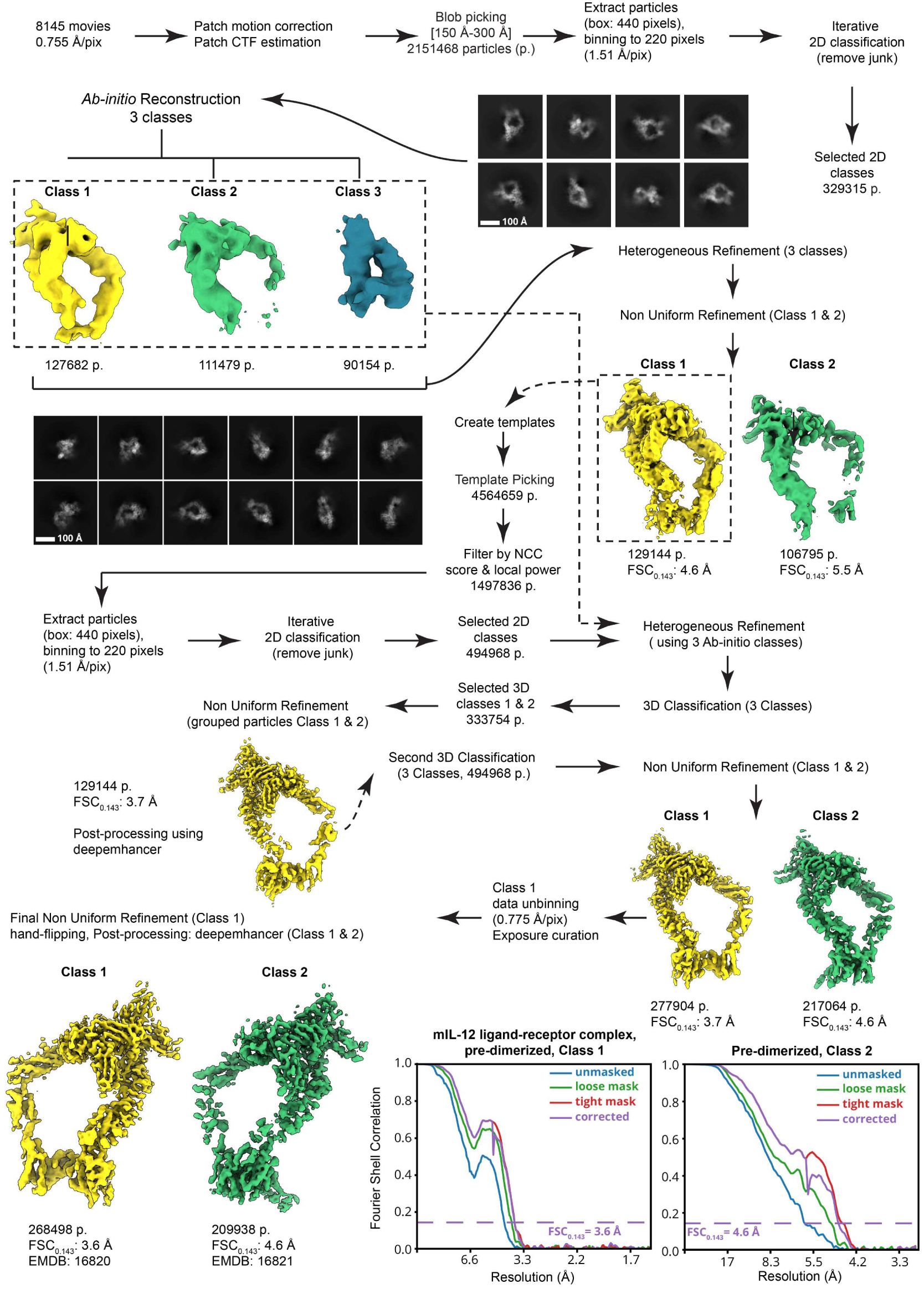
Cryo-EM data processing workflow for the pre-dimerized mIL-12:mIL-12Rβ1_D1-D5_-DAPK1_302-330_:mIL-12Rβ2_D1-D6_-CaM complex. Data processing was performed in CryoSPARC v3.3.2^24,25^, and map post-processing was performed using DeepEMhancer^27^. Gold-standard Fourier Shell Correlation (FSC) curves are shown after applying either no mask (blue), a loose mask (green), or a tight mask (red) to both half maps before calculating the FSC. The corrected FSC (purple) is calculated using the tight mask with correction by noise substitution^41^. The estimated resolution at FSC = 0.143 (dotted purple lines) is shown for the corrected FSC curves (purple lines).

**Extended Data Figure 3:**
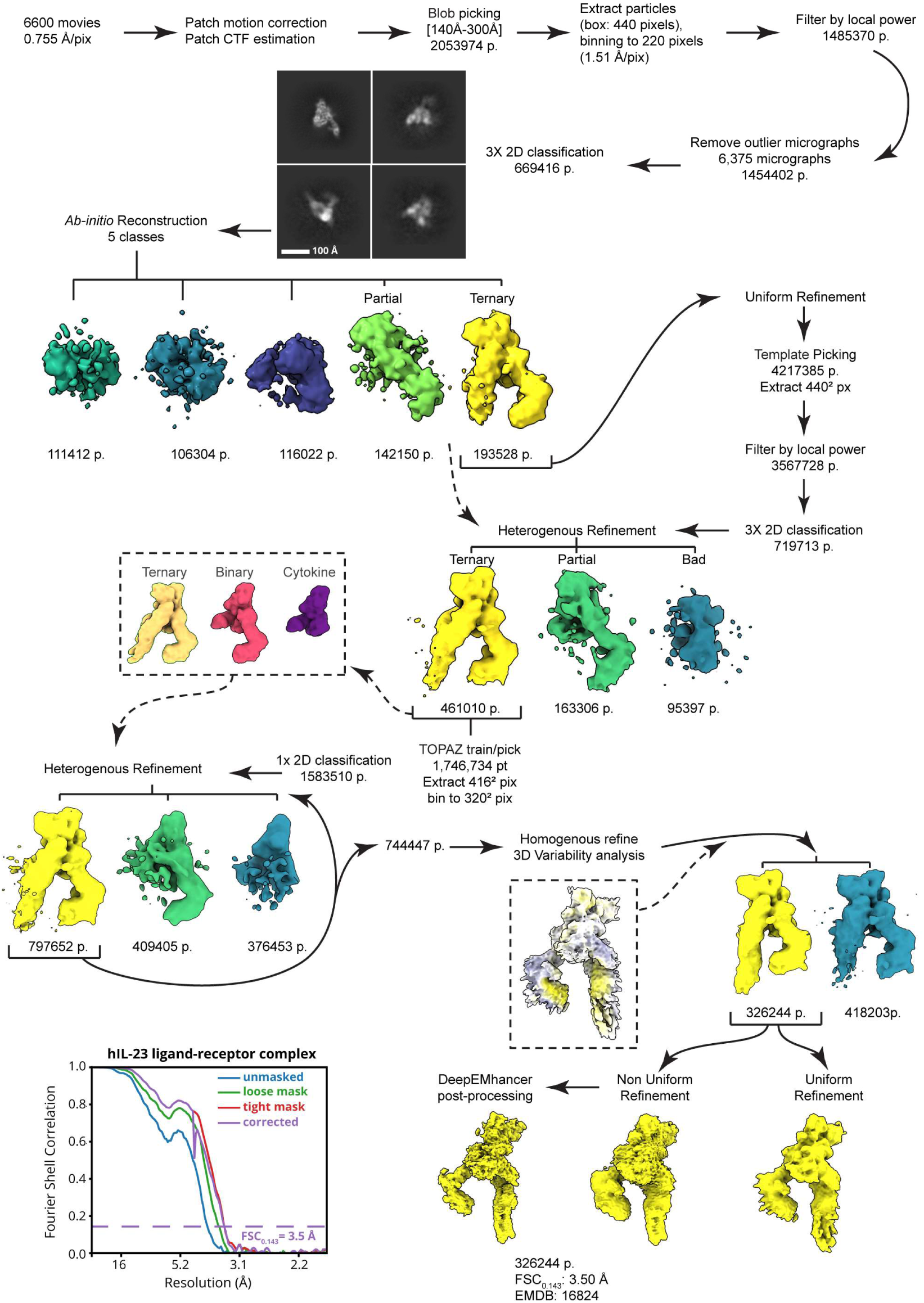
Cryo-EM data processing workflow for the pre-dimerized hIL-23:hIL-12Rβ1_D1-D5_-DAPK1_302-330_:hIL-23R-CaM complex. Data processing was performed in CryoSPARC v3.3.2^24,25^, and map post-processing was performed using DeepEMhancer^27^. Gold-standard Fourier Shell Correlation (FSC) curves are shown after applying either no mask (blue), a loose mask (green), or a tight mask (red) to both half maps before calculating the FSC. The corrected FSC (purple) is calculated using the tight mask with correction by noise substitution^41^. The estimated resolution at FSC = 0.143 (dotted purple lines) is shown for the corrected FSC curves (purple lines).

**Extended Data Figure 4:**
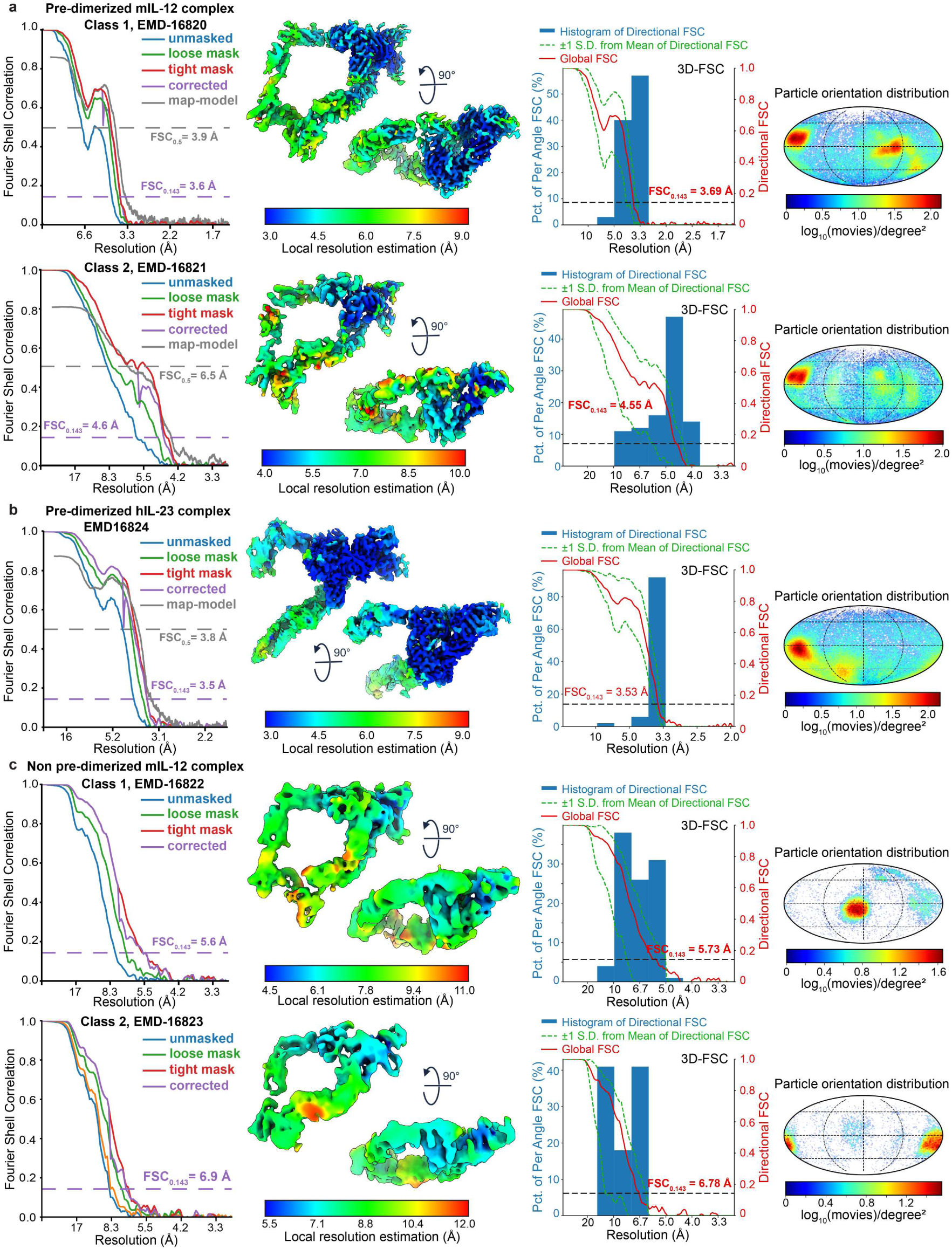
Gold-standard Fourier Shell Correlation (FSC) curves, local resolution estimation and 3D-FSC analysis of reported maps: **a - c**, Gold-standard FSC curves for Class 1 & 2 (a) of the pre-dimerized mIL-12:mIL-12Rβ1_D1-D5_-DAPK1_302-330_:mIL-12Rβ2_D1-D6_-CaM complex, the pre-dimerized hIL-23:hIL-12Rβ1_D1-D5_-DAPK1_302-330_:hIL-23R-CaM complex (b), and Class 1 & 2 (c) of the non pre-dimerized mIL-12:mIL-12Rβ1_D1-D5_-Strep-II:mIL-12Rβ2_D1-D6_-Strep-II complex. FSC curves are calculated after applying either no mask (blue), a loose mask (green), or a tight mask (red) to both half maps. The corrected FSC (purple) is calculated using the tight mask with correction by noise substitution^41^. The estimated resolution at FSC = 0.143 (dotted purple lines) is shown for the corrected FSC curves (purple lines). Map-to-model FSC curves are shown in grey, along with the estimated resolution at FSC = 0.5 (dotted grey lines). To the right of each set of FSC curves, a local resolution coloring of the corresponding 3D reconstruction is displayed, along with 3D-FSC plots calculated using the Remote 3D-FSC Processing Server^42^ and particle orientation distribution plots generated using an adapted script from cryoEF v1.1.0^43^.

**Extended Data Figure 5:**
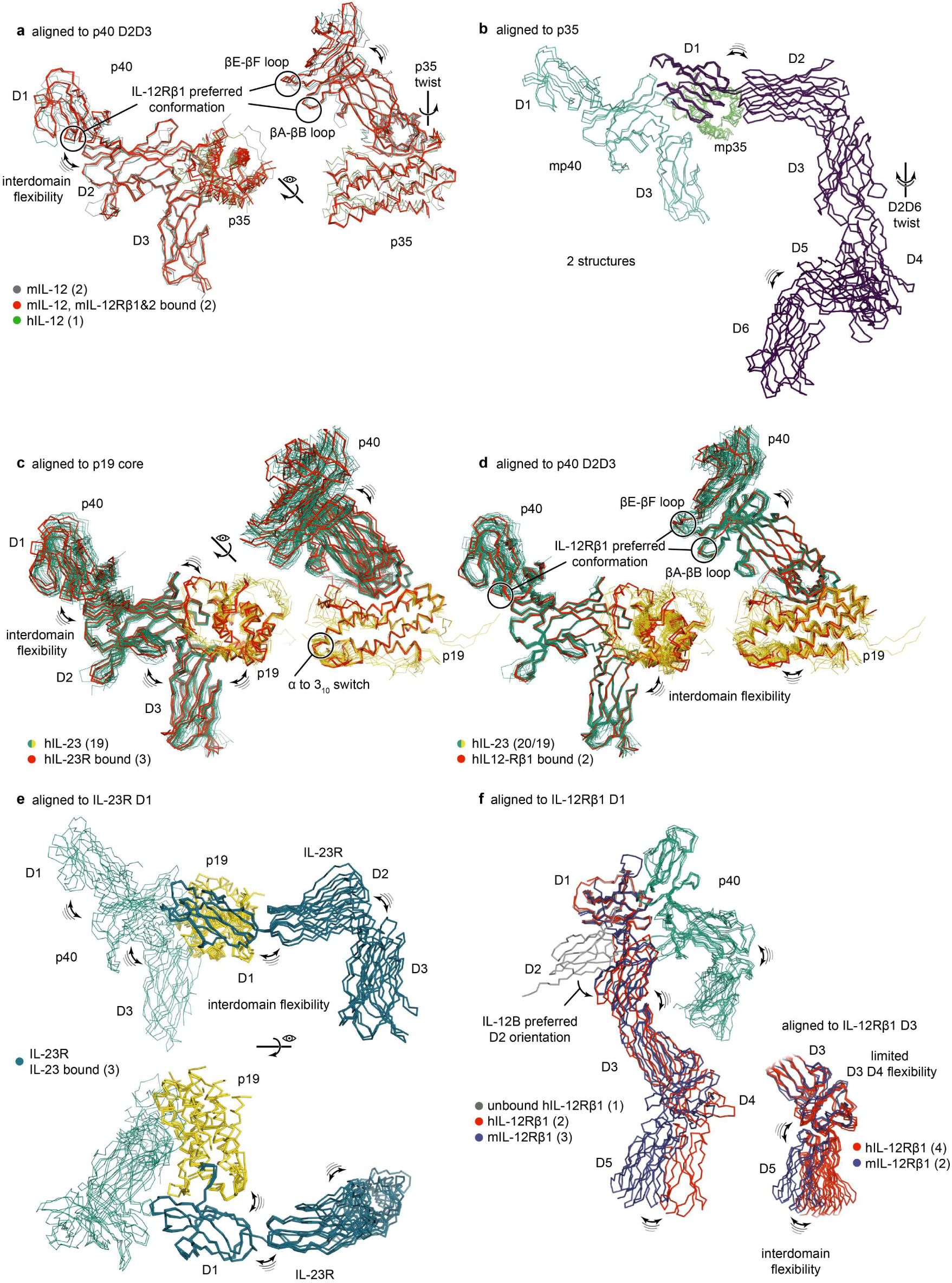
Experimentally observed flexibility in the IL-12 and IL-23 protein complexes. **a)** structural superposition of hIL-23 (21 observations total) using the p19 helices as a reference showing inter domain flexibility. IL-23R bound structures display an α-helical to 3_10_-helical switch at the N-terminal tip of the D-helix which is the interface hotspot. **b)** structural superposition of hIL-23 (21 observations of hIL-23 and 1 hp40 only) using the p40_D2D3_ as a reference showing inter domain flexibility. hIL-12Rβ1 bound structures display a preferred orientation of the p40_D1_ especially with regards to the p40_D1_βA-βB and βE-βF loops which are part of the interface. **c)** Structural superposition of IL-12 (5 observations) using p40_D2D3_ as a reference showing inter domain flexibility. mIL-12Rβ1_D1-D5_ bound structures display a preferred orientation of the p40_D1_ especially with regards to the p40_D1_βA-βB and βE-βF loops which are part of the interface. The p35 subunit is also twisted away from p40 upon mIL-12Rβ2 binding. **d)** Structural superposition of mIL-12Rβ2 (2 structures) using p35 as a reference displays flexibility beyond the p35: mIL-12Rβ2_D1_ interface. **e)** Structural superposition of hIL-23R using hIL-23R_D1_ as reference displays flexibility in the receptor domains a well as in cytokine binding. Models utilized in figures are PDB IDs: 1f45, 3hmx, 3duh, 3d85, 3d87, 3qwr, 4grw, 5mj3, 5mj4, 5mxa, 5mzv, 5njd, 6uib, 6wdq, 6sff, 6smc, 6sp3, 7pur, 7r3n, 8odz, 8oe0, 8oe4, 8cr6, 8cr5, 8cr8, 8c7m, 8odx.

**Extended Data Figure 6:**
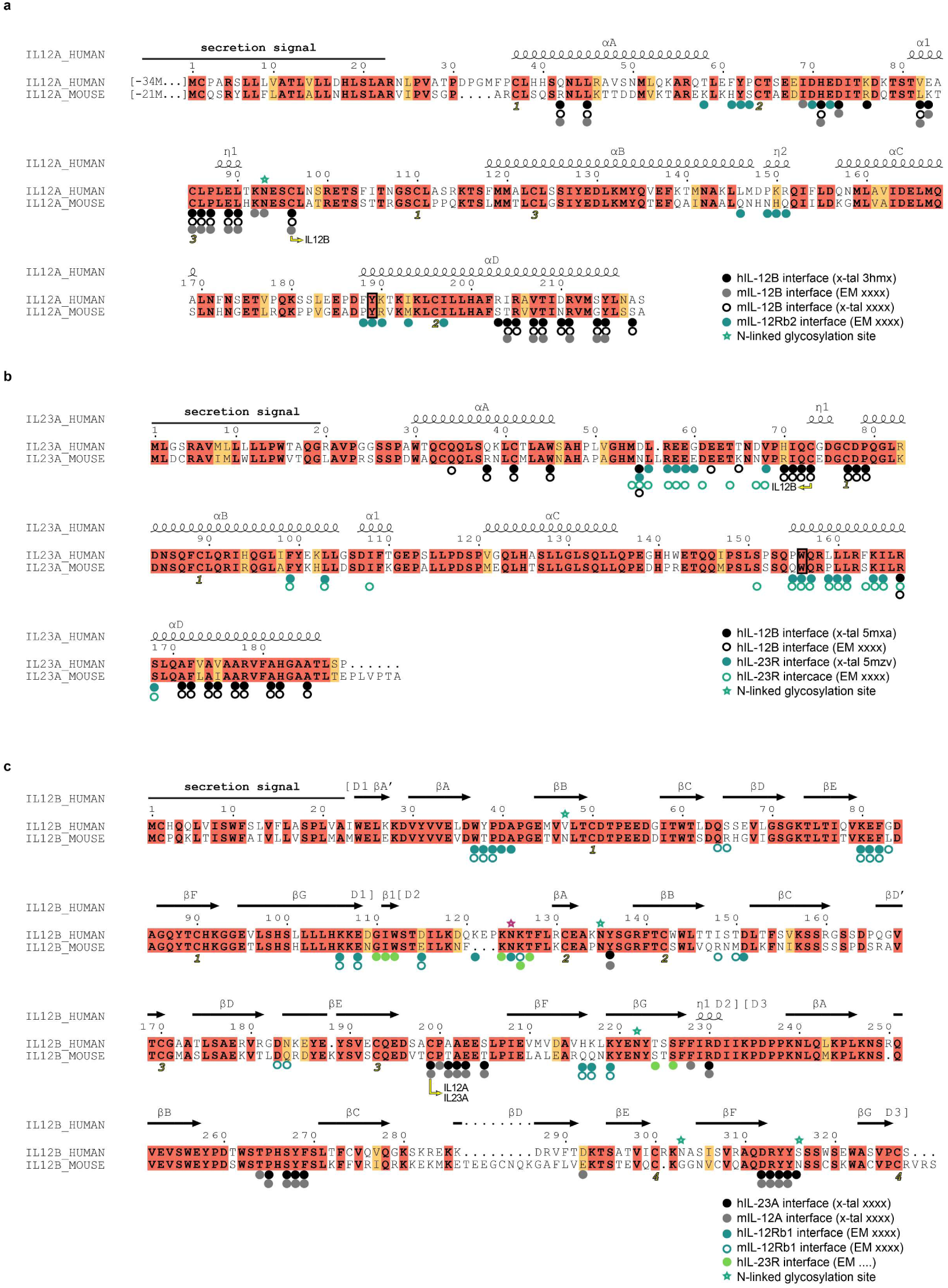
Pairwise sequence alignments of human and murine IL-12A, IL-23A and IL-12B. Sequence alignments are shown between human and murine orthologs of IL-12A (a), IL23A (b) and IL-12B (c). Secretion signals are indicated with a black line, and secondary structure elements are indicated with helices and arrows for α-helices and β-strands respectively. Residues involved in specific interaction interfaces are annotated using colored dots, and N-linked glycosylation sites are annotated using a star. Pairwise alignments were performed using MAFFT^44^, and ESPript^45^ was used for display.

**Extended Data Figure 7:**
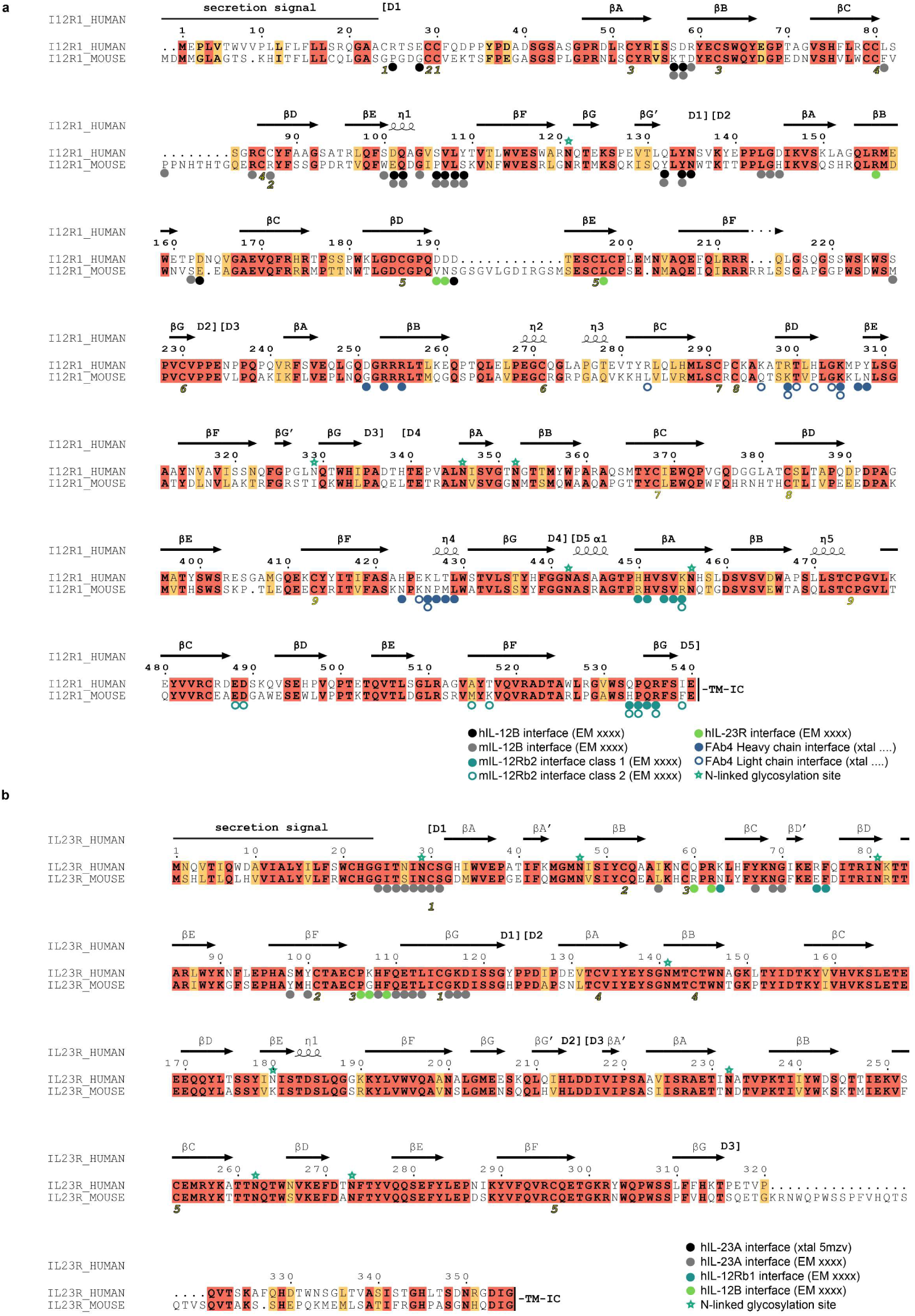
Pairwise sequence alignments of human and murine IL-12Rβ1 and IL-23R. Sequence alignments are shown between human and murine orthologs of IL-12Rβ1 (a) and IL-23R (b). Secretion signals are indicated with a black line, and secondary structure elements are indicated with helices and arrows for α-helices and β-strands respectively. Residues involved in specific interaction interfaces are annotated using colored dots, and N-linked glycosylation sites are annotated using a star. Pairwise alignments were performed using MAFFT^44^, and ESPript^45^ was used for display.

**Extended Data Figure 8:**
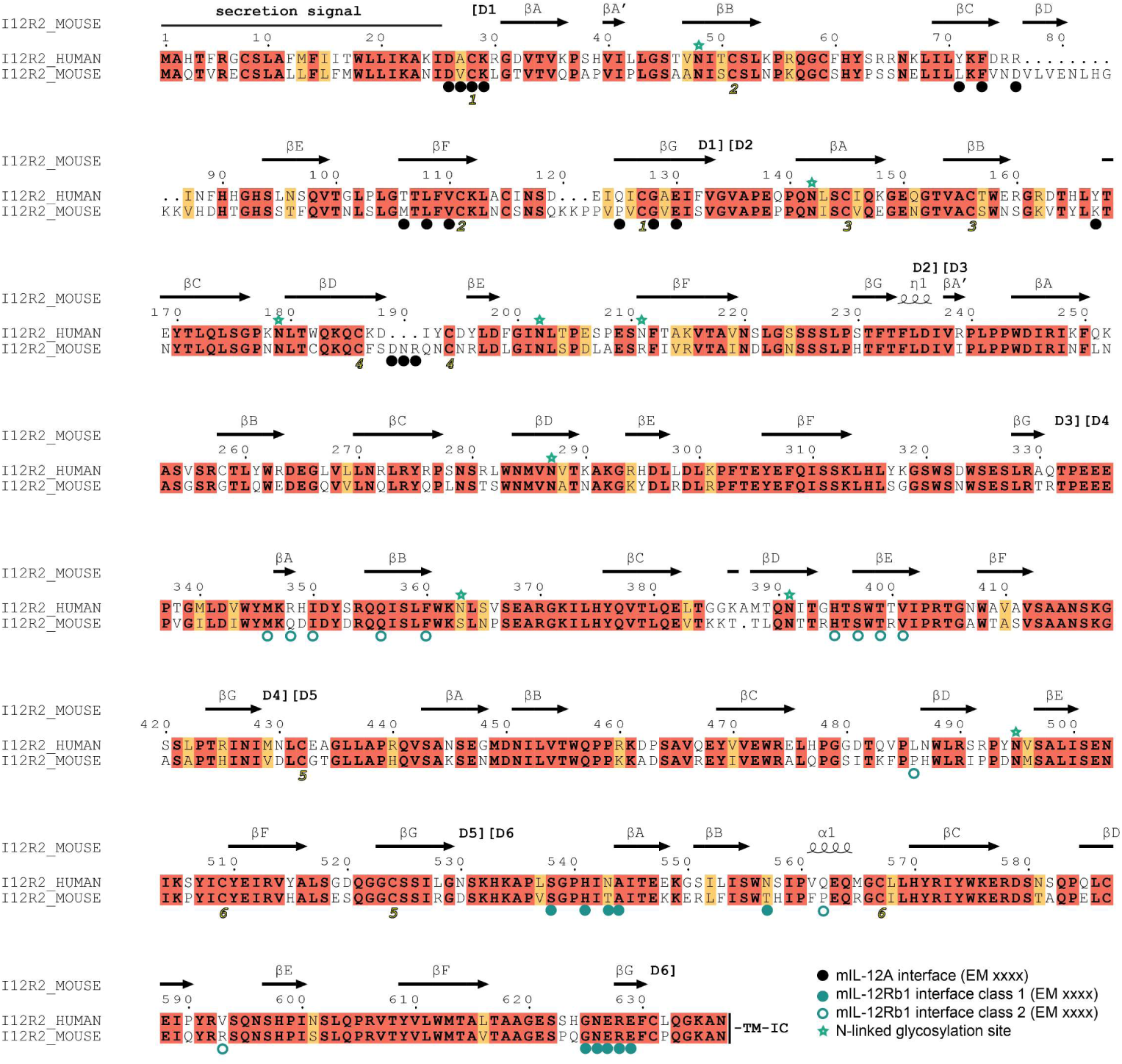
Sequence alignment of human and murine IL-12Rβ2. A sequence alignment is shown between human and murine orthologs of IL-12Rβ2. Secretion signals are indicated with a black line, and secondary structure elements are indicated with helices and arrows for α-helices and β-strands respectively. Residues involved in specific interaction interfaces are annotated using colored dots, and N-linked glycosylation sites are annotated using a star. Pairwise alignments were performed using MAFFT^44^, and ESPript^45^ was used for display.

**Extended Data Figure 9:**
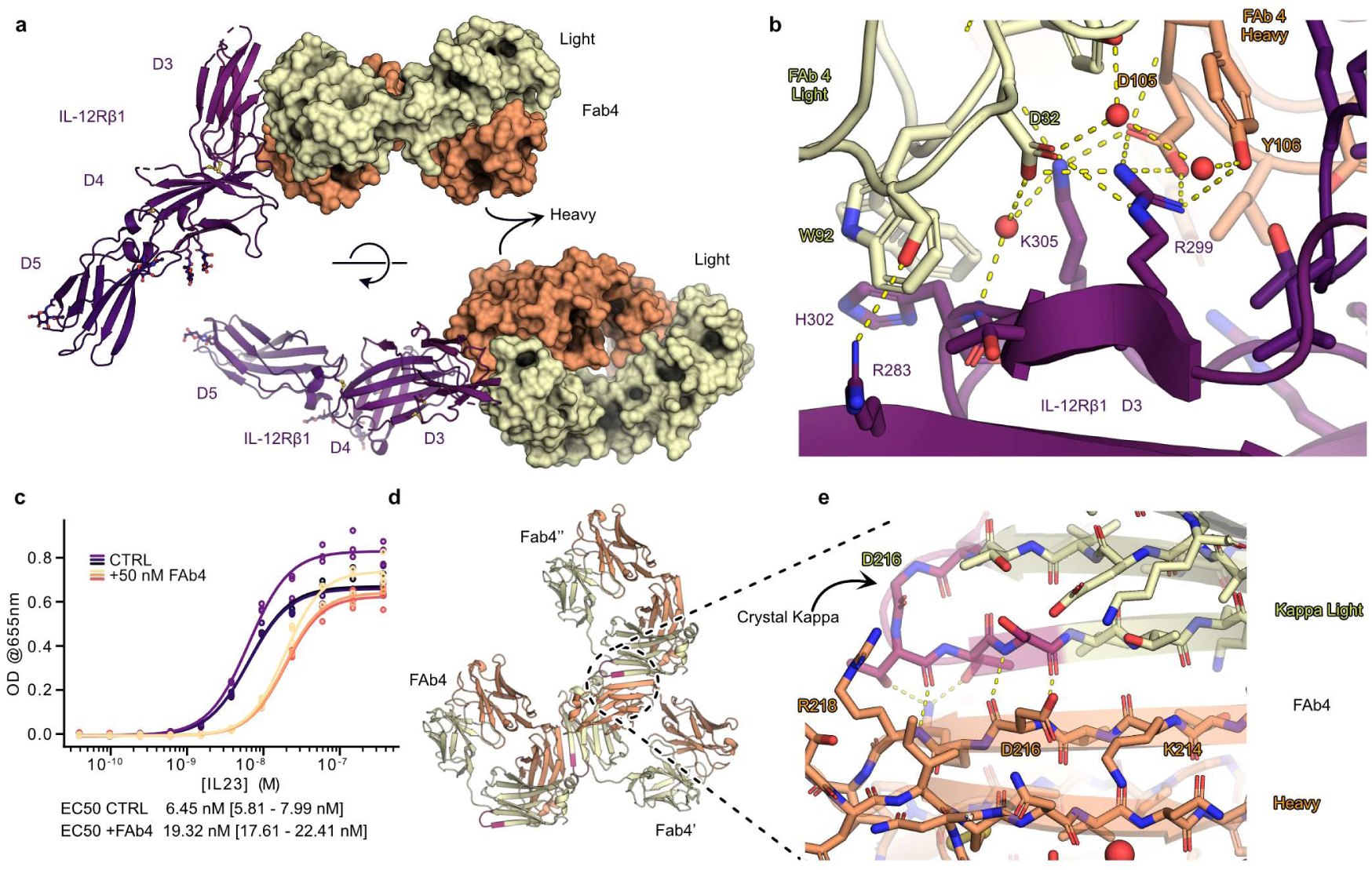
Details of the IL-12Rβ1_D3-D5_:Fab4 interaction. A) Cartoon representation of the IL-12Rβ1_D3-D5_:Fab4 interaction. Receptor fragment in cartoon representation and Fab4 chain in surface representation. B) Detailed view of the IL-12Rβ1_D3-D5_:Fab4 interaction around residue R299 including some waters (red spheres) which are part of the interface. C) Dose-response curve of a SEAP based IL-23 reporter cellular assay in presence or absence of Fab4. EC_50_ values are reported together with their 95% confidence interval. 3 biological replicates, each in triplicate are shown. D) Cartoon representation of the crystal packing interaction of the better defined (lower B-factor) Fab4 copy interacting with two crystallographic copies (IL-12Rβ1 fragment and other complex not shown). E) The engineered Crystal Kappa mutation (magenta) near the C-terminus of the Light chain (wheat) forms an anti-parallel beta-sheet with the C-terminal beta strand of the Heavy chain (orange) of the neighboring Fab fragment allowing for a crystal lattice.

**Supplementary Figure 1:**
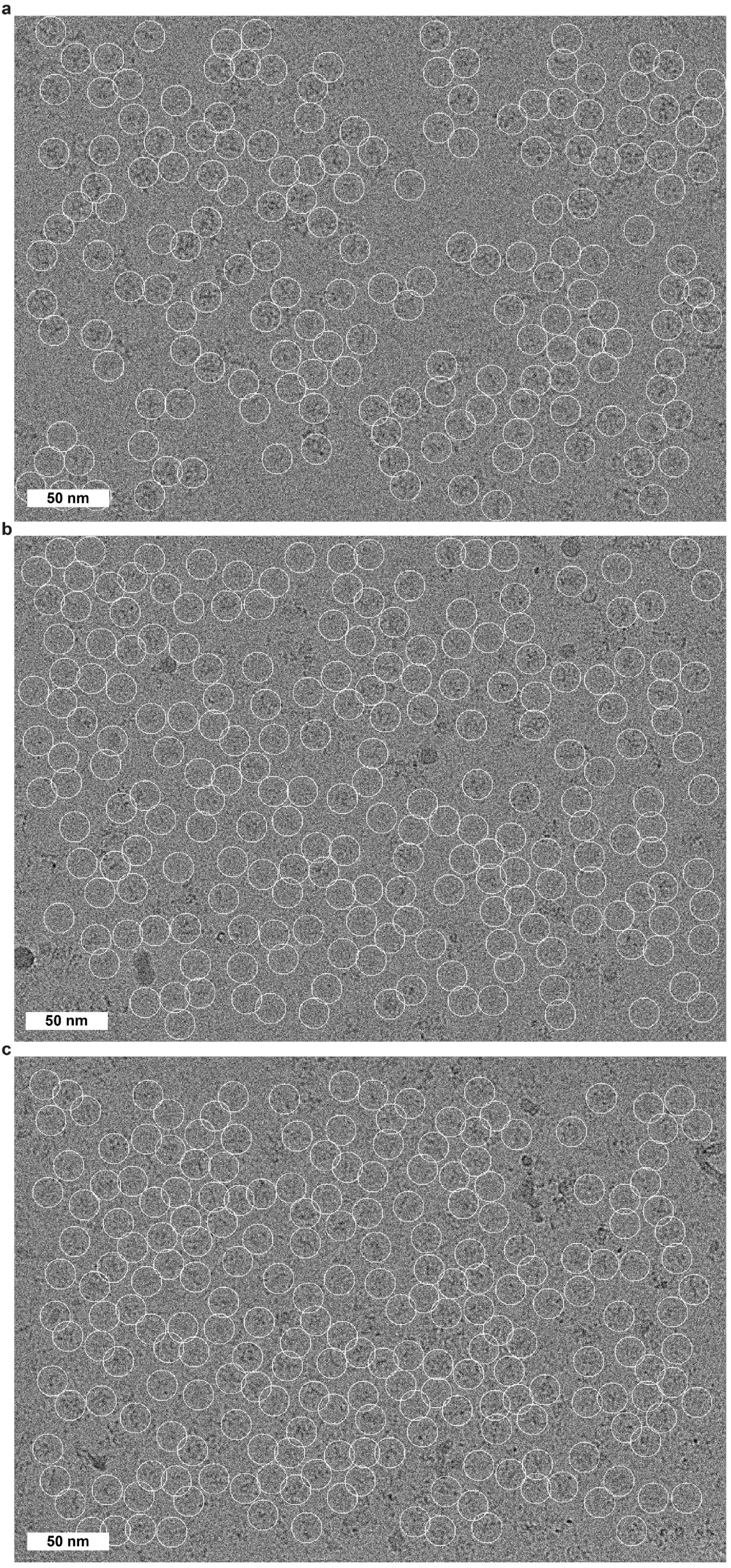
Motion corrected and CTF estimated micrographs. Motion corrected and CTF estimated micrographs are displayed of pre-dimerized mIL-12 (a), non pre-dimerized mIL-12 (b) and hIL-23 (c) ligand-receptor assemblies. Picked particles are encircled and a 50 nm scale bar is shown on the bottom left of each micrograph.

**Supplementary Table 1.** X-ray crystallographic data collection and refinement statistics.

|  | mIL-12<br>(PDB 8cr6) | mIL-12B<sub>C197S</sub>:<br>mIL-12Rβ1<sub>D1-D2</sub><br>complex<br>(PDB 8cr5) | hIL-23<br>(PDB 8cr8) | hIL-12Rβ1<sub>D3-D5</sub>:<br>Fab4<sub>CrystallKappa</sub><br>complex<br>(PDB 8c7m) | hIL-12Rβ1<sub>D3-D5</sub>:<br>Fab4:anti-Kappa-VHH complex<br>(PDB 8odx) |
| --- | --- | --- | --- | --- | --- |
| **Data collection** |  |  |  |  |  |
| Space group | *P* 2<sub>1</sub> 2<sub>1</sub> 2<sub>1</sub> | *P* 1 2<sub>1</sub> 1 | *P* 1 2<sub>1</sub> 1 | *P* 2<sub>1</sub> 2<sub>1</sub> 2<sub>1</sub> | *P* 4<sub>1</sub> 3 2 |
| Cell dimensions<br>*a*, *b*, *c* (Å) | 59.11, 70.77, 127.37 | 29.22, 165.09, 51.87 | 67.75, 59.84,<br>69.99 | 73.66 140.18<br>219.36 | 231.40, 231.40,<br>231.40 |
| $\alpha$, $\beta$, $\gamma$ (°) | 90.00, 90.00, 90.00 | 90.00, 90.44, 90.00 | 90.00, 98.06,<br>90.00 | 90.00, 90.00,<br>90.00 | 90.00, 90.00,<br>90.00 |
| Resolution (Å) | 47.34 - 2.85<br>(2.95 - 2.85)* | 43.92 - 2.15<br>(2.23 - 2.15)* | 69.30 - 2.00<br>(2.12 - 2.00) | 140.18 - 2.56<br>(2.71 - 2.56) | 231.40 - 4.40<br>(4.67 - 4.40) |
| *R*<sub>meas</sub> (%) | 5.7 (132.8) | 17.2 (127.8) | 7.3 (141.1) | 10.4 (208.3) | 23.1 (394.1) |
| *<I / σI>* | 13.85 (1.07) | 7.18 (1.07) | 13.90 (1.02) | 19.16 (1.30) | 18.54 (1.36) |
| Completeness (%) | 96.20 (96.66) | 99.36 (99.33) | 94.3 (74.1) | 99.8 (98.9) | 100.0 (100.0) |
| Redundancy | 2.9 (2.9) | 3.6 (3.7) | 6.58 (5.07) | 13.60 (13.28) | 70.1 (71.3) |
| Wilson B factor (Å<sup>2</sup>) | 97.92 | 38.28 | 55.53 | 76.27 | 224.70 |
| **Refinement** |  |  |  |  |  |
| Resolution (Å) | 2.85 | 2.15 | 2.00 (2.05-2.00) | 2.56 (2.62 -<br>2.56) | 4.40 (4.56-4.40) |
| No. unique reflections | 12528 (1245) | 26465 (2664) | 35506 (1809) | 73930 (5095) | 13964 (1363) |
| *R*<sub>work</sub> /<br>*R*<sub>free</sub> (%) | 25.46 (49.48)/<br>29.59 (62.42) | 23.96 (38.50)/<br>28.20 (41.34) | 23.50<br>(38.04)/25.59<br>(37.06) | 20.34<br>(35.46)/22.82<br>(38.38) | 27.96(44.86)/<br>31.15(45.73) |
| No. atoms |  |  |  |  |  |
| Protein | 3584 | 3186 | 3438 | 10815 | 6382 |
| Ligand/ion | 53 | 64 | 73 | 72 | N.A. |
| Water | 2 | 114 | 84 | 24 | N.A. |
| *B*-factors (Å<sup>2</sup>) |  |  |  |  |  |
| Protein | 110.42 | 66.28 | 65.43 | 104.24 | 270.41 |
| Ligand/ion | 121.29 | 59.26 | 82.01 | 105.20 | N.A. |
| Water | 63.05 | 48.74 | 55.38 | 67.03 | N.A. |
| R.m.s. deviations |  |  |  |  |  |
| Bond lengths (Å) | 0.002 | 0.004 | 0.003 | 0.002 | 0.004 |
| Bond angles (°) | 0.490 | 0.966 | 0.525 | 0.56 | 0.824 |

\*Values in parentheses correspond to the highest-resolution shell. Ramachandran statistics can be found in the Methods section under 'X-Ray crystallographic Model Building and Refinement'.

**Supplementary Table 2.** Summary of cloned constructs.

| Protein | Database ref.<br>(UniProt/PDB) | Delineation | Mutations | Vector | Plasmid ID<br>(BCCM)<sup>‡‡</sup> |
| --- | --- | --- | --- | --- | --- |
| **hIL-23A** | Q9NPF7 | <sup>†</sup>GauLucSS-[S27-P189]-Casp<sup>‡</sup>-His<sub>6</sub><br>T2A |  | pHLsec | **LMBP 13691** |
|  |  | mIgHVSS-[SRN]-[V22-P189]-His<sub>6</sub> |  |  | **LMBP 13692** |
| **hIL-12B** | P29460 | <sup>†</sup> [M1-S328]<br>GauLucSS-[LE]-[I23-S328] | N303D | pHLsec | **LMBP 13693** |
| **hIL-23R** | Q5VWK5 | [M1-D353]- Calmodulin |  | pHLsec | **LMBP 13694** |
| **hIL-12Rβ1<sub>D1-D5</sub>** | P42701 | [M1-E540]- DAPK1 |  | pHLsec | **LMBP 13695** |
| **hIL-12Rβ1<sub>D3-D5</sub>** | P42701 | RTPμSS-[P232-E540]-Casp-AviTag-His<sub>6</sub> |  | pHLsec | **LMBP 13696** |
| **mIL-12A** | P43431 | [M1-A215]-Casp-AviTag-His<sub>6</sub> |  | pHLsec | **LMBP 13683** |
| **mIL-12B** | P43432 | [M1-S335] |  | pHLsec | **LMBP 13682** |
| **mIL-12B<sub>C197S</sub>** | P43432 | [M1-S335]-His<sub>6</sub> | C197S | pHLsec | **LMBP 13684** |
| **mIL-12Rβ1<sub>D1-D5</sub>** | Q60837 | GauLucSS-[Q20-E561]-DAPK1 |  | pHLsec | **LMBP 13685** |
|  |  | GauLucSS-[Q20-E561]-Strep-II Tag |  |  | **LMBP 13686** |
| **mIL-12Rβ1<sub>D1-D2</sub>** | Q60837 | GauLucSS-[Q20-L258]-His<sub>6</sub> |  | pHLsec | **LMBP 13687** |
|  |  | GauLucSS-[Q20-L258]-Casp-AviTag-His<sub>6</sub> |  |  | **LMBP 13688** |
| **mIL-12Rβ2<sub>D1-D6</sub>** | P97378 | GauLucSS-[N24-N637]-Calmodulin |  | pHLsec | **LMBP 13689** |
|  |  | GauLucSS-[N24-N637]-Strep-II Tag |  |  | **LMBP 13690** |
| **Calmodulin tag** | P0DP23 | GTGGSGGSGG-[L5-K149] |  |  |  |
| **DAPK1 tag** | P53355 | GTGGSGGSGG-[A300-S319] |  |  |  |
| **aKappa-VHH** | PDB 6ww2,<br>chain K | PelB-6His-Casp-[] |  | pEt15b | **LMBP 13697** |
| **Fab4 Heavy** | US8574573 n°58 | mIgHeavySS-[]-Casp-6His |  | pHLsec | **LMBP 13698** |
| **Fab4 Light** | US8574573 n°70 | GauLucSS-[] |  | pHLsec | **LMBP 13699** |
| **Fab4**<br>**Light<sub>CrystallKappa</sub>** | US8574573 n°70 | GauLucSS-[] | H215del,<br>L218T,<br>S219T,<br>P221del | pHLsec | **LMBP 13700** |

<sup>†</sup> hIL-23A and hIL-12B were cloned into a contiguous gene separated by a T2A peptide.
<sup>‡</sup> The Caspase3 protease site (Casp) is encoded by the DEVD/G peptide. GauLucSS = *Gaussia princeps* luciferase secretion signal, RTPμSS = chicken (*Gallus gallus*) Receptor Tyrosine Phosphatase (RTP) μ-like secretion signal.
<sup>‡‡</sup> IDs can be entered at https://bccm.belspo.be/catalogues/genecorner-plasmids-catalogue-search to find the relevant deposited plasmid.

